# Chemical Mixtures in River Basins: Combining Additive and Independent Effects with Probabilistic Risk Modelling

**DOI:** 10.64898/2026.06.03.729784

**Authors:** S. Jannicke Moe, Anders L. Madsen, Sophie Mentzel, Karel Viaene, Karel Vlaeminck, Merete Grung, Samantha Martins, Gašper Šubelj, Sam Welch, Frederik Verdonck

## Abstract

Chemical mixtures and potential cocktail effects in aquatic ecosystems are recognised as a threat for river basins world-wide. Probabilistic risk approaches are becoming more common in environmental risk assessment, and offer new opportunities for metodological challenges such as of mixture risk characterisation. The Concentration Addition (CA) concept is commonly used in lower-tier risk assessment (e.g., sum of risk quotients), as a pragmatic and protective method. However, the alternative Independent Action (IA) concept can easily be implemented in probabilistic risk calculation (e.g., joint probability of threshold exceedances).

We have developed a multi-level probabilistic model for integrating these two concepts, formulated as an object-oriented Bayesian network (BN). First, probabilistic risk quotients (RQ) are calculated for individual substances, as probability distributions of environmental concentrations divided by a threshold environmental value. Next, the CA concept is applied within groups of substances by summing the RQ distributions. Finally, the IA concept is applied across the different substance groups, assuming independent modes of action, to combine RQ distributions by joint probability calculation (“OR” expressions). Predicted exposure concentrations were obtained from the ENCORE fate model, a process-based model for simulation of chemicals in river basins across Europe.

Here we present a pilot study focusing on a subset of the substances (15 pesticides) and river basins (in Belgium), as a proof-of-concept. The purpose of this pilot study was to demonstrate a novel probabilistic approach to mixture risk characterisation, by combining the CA and IA concepts in a multi-level BN. The results were consistent across scenarios as well as with literature, with CA-based risk characterisations being slightly higher the IA-based. The combined CA+IA-based risk represents a reasonable compromise. Sensitivity analysis of the BN can provide an effective ranking of the risk-driving substances and groups, to support chemical prioritisation and risk managment.

**Key points:**

1. A multi-level Bayesian network (BN) was developed for probabilistic calculation of environmental risk from chemical mixtures, based on risk quotients (RQ) calculated for individual substances, with a selection of 15 pesticides in Belgium as a pilot study.
2. Predicted environmental concentrations (PEC) are obtained from the ENCORE exposure model; a process-based model which can simulate transport, fate and concentration of >1000 substances (pesticides, pharmaceuticals, etc.) in rivers subcatchments across Europe based on chemical use and emission.
3. The BN’s risk calculation uses substance groups (Fungicides, Herbicides and Insecticides) to combine two classical mixture concepts: (1) Concentration Addition: sum of RQs within groups, followed by (2) Independent Action: joint probability of threshold exceedance for one or more groups.
4. The resulting rankings of risk-driving substances by the BN for this pilot study are robust across scenarios, suggesting a potential for expanding this generic BN approach to different mixtures with higher numbers of substances and groups, and to larger regions of Europe.

## 1. Introduction

Chemical mixtures are ubiquitous in aquatic ecosystems (EEA 2018). Environmental risk assessment of chemical mixtures is challenging because of the multitude of possible combinations that may occur (Holmes et al., 2017). While individual substances can have concentrations below the regulatory limit (e.g. Environmental Quality Standards in Europe), the combination of substances can still be harmful to an aquatic ecosystem. It has been argued that ecological impacts of mixtures is not sufficiently considered in the current chemical regulation under REACH (Registration, Evaluation, Authorization and Restriction of Chemicals) in the EU (Backhaus et al., 2025). More attention therefore is needed to address the risk posed by the ‘cocktail effect’ of lower concentrations of chemicals in European lakes, rivers and other surface water bodies (EEA 2018).

While mixture toxicity can be determined experimentally for a limited number of substances, suitable models can predict the combined effects of mixtures more efficiently. Environmental risk assessment (ERA) for chemical mixtures at regional scales therefore relies on predictive models for characterisation of the combined risk to ecological assessment endpoints (Bopp et al., 2015). However, methodological approaches to mixture risk assessment have been subject to much debate. Established methods for cumulative risk assessment are usually based on simple approximations referred to as concentration addition (CA) (Sigurnjak Bureš et al., 2021)). These can involve summation of scaled concentration metrics referred to as toxic units (Backhaus et al., 2013), risk quotients (Duarte et al., 2022) or hazard indexes (Rodea-Palomares et al., 2023). An alternative method, referred to as independent action (IA), can be used for calculating mixture risk as the joint probability of specified effect for each substance (e.g., Neelamraju et al., 2025). Other studies have shown that a combination of the CA- and IA-based methods may be useful (Sigurnjak Bureš et al., 2021). In the EU, many scientists have advocated that chemical management under REACH should incorporate a mixture allocation factor (MAF) for better management of chemical “cocktails” (Backhaus et al., 2025). In contrast, some authors have stated that mixture risk in European surface waters is dominated by a low number of substances, which does not support the introduction of a MAF (Rodea-Palomares et al., 2023). Others have argued that computational approaches within the current ERA paradigm are too reductionist, although acknowledging the need for pragmatic solutions (Topping et al., 2026). All in all, the ongoing debates illustrate the need for continued development and evaluation of modelling methodologies for mixture risk assessment.

Probabilistic risk modelling is becoming more common in human and ecological risk assessment (Maertens et al., 2022), and can offer new approaches to fundamental challenges such as the characterisation of risk from chemical mixtures. In this study, we explore the use of Bayesian Network (BN) modelling as a tool to support probabilistic risk assessment of mixtures. The probabilistic risk characterisation is based on risk quotients (RQs) quantified by discrete probability distributions, in the same manner as described by for example (Carriger and Barron, 2020; Mentzel et al., 2022; Welch et al., 2024). In this way, the spatio-temporal variability and other sources of uncertainties can be characterised. BN methodology has gained popularity as a tool for environmental risk assessment during the last decade (Moe et al., 2021). Moreover, probabilistic risk calculation offers novel alternatives for characterisation of mixture risk. While CA-based methods are currently the most used in risk assessment of mixtures, the IA concept lends itself more easily to probabilistic risk calculation assuming independent events. Here, we explore a combination of the two methods (CA+IA) by a two-step calculation in probabilistic framework (Table 1): First, cumulative effects within substance groups are summed according to the CA principle; secondly, the combined risk across substance groups is calculated as the joint probability of independent events according to the IA principle. The procedure for combining CA and IA calculations is comparable to the two-stage prediction model presented by Junghaus (2004), as well as the two-step mixed-model for single species or species assemblages introduced by (de Zwart and Posthuma, 2005). In our study, the combined CA+IA approach is further generalised by using RQs as a simplified risk metric. Furthermore, the approach is implemented in BN methodology for effective probabilistic risk modelling with a graphical layout.

**Table 1.**
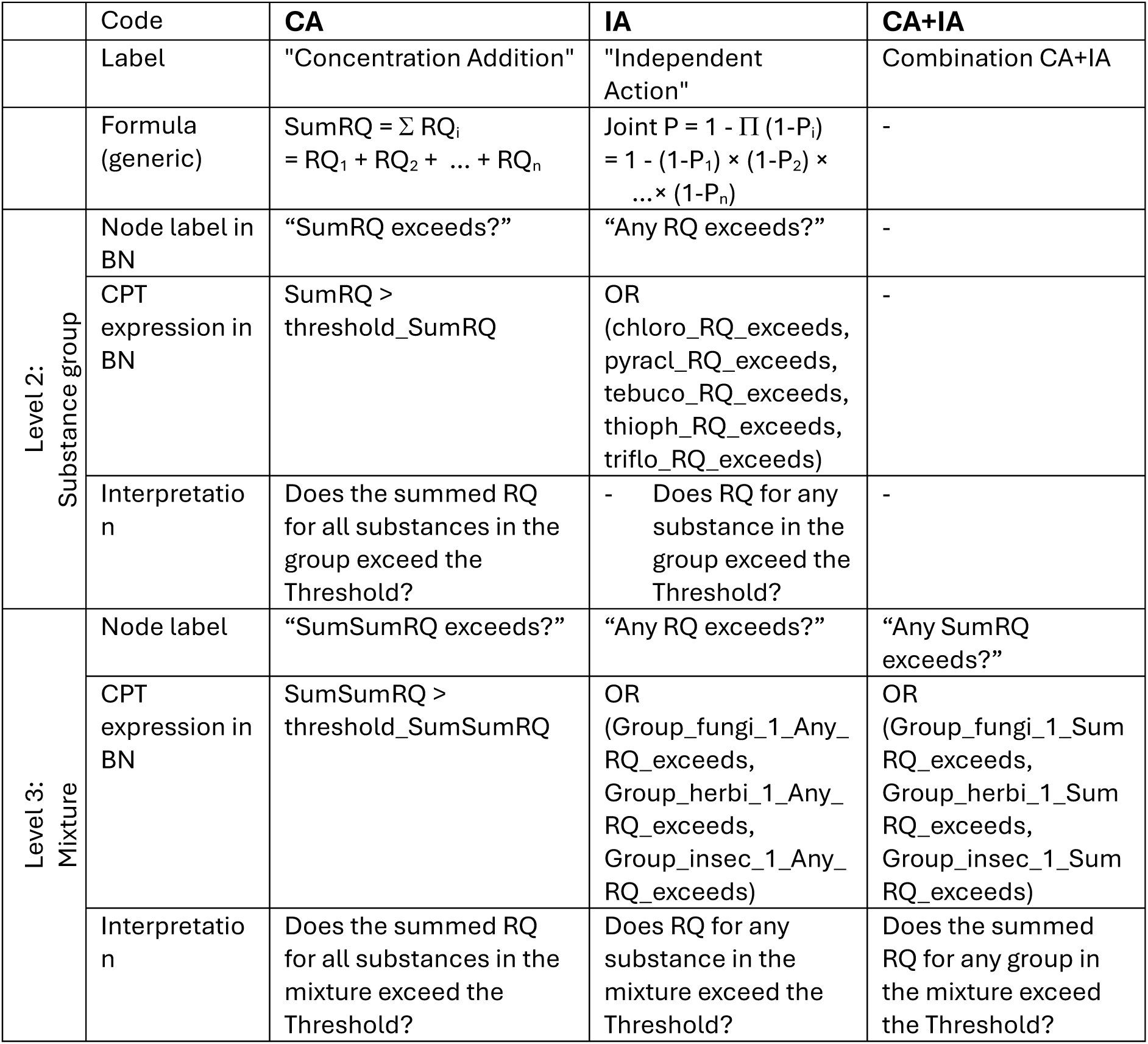
The three methods for mixture risk calculation applied in this study. CPT = conditional probability table, RQ = risk quotient.

To enable the risk characterisation across multiple levels (substances, groups and mixture) in a systematic and traceable way, we have designed a hierarchical probabilistic model and constructed this as a multi-level object-oriented Bayesian network (OOBN). The OOBN methodology makes it possible to trace and display all calculations involved in all three mixture risk calculation methods (CA, IA and CA+IA) across the three levels, through modular submodels. This novel modelling approach to mixture risk is applied to pilot study comprising a selection of pesticides of concern for aquatic ecosystems in Belgium (Figure 1, Table 2).

**Figure 1.**
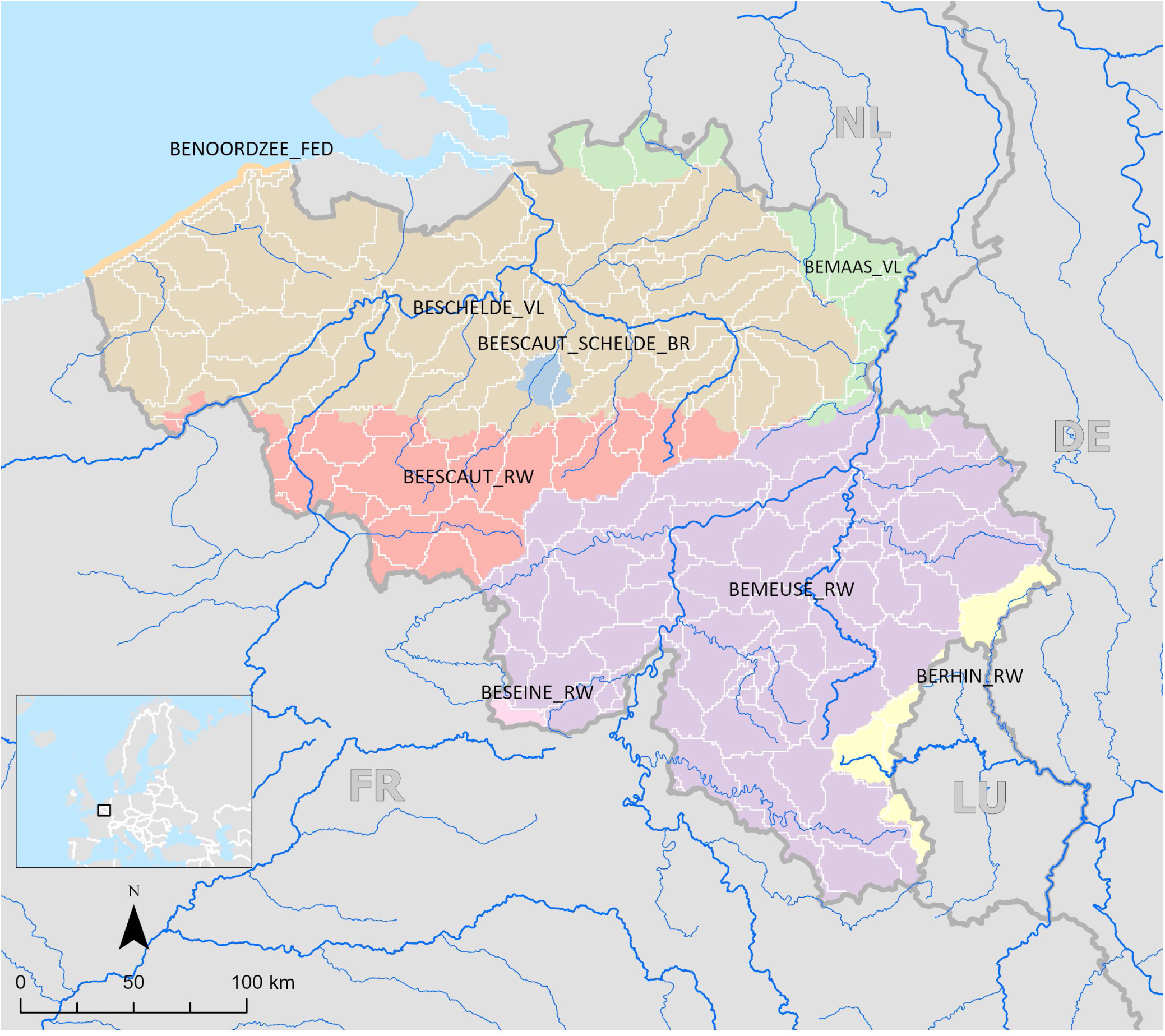
Map of Belgium showing the spatial structures of the pilot study: subcatchments (spatial units of exposure model, white outlines) and river basin disctricts (RBD, spatial units of the Bayesian network, different colours). The RBDs included in the BN represent the rivers Meuse (BEMEUSE_RW, BEMAAS_VL), Scheldt (BEESCAUT_RW, BESCHELDE_VL), and Rhine (BERHIN_RW). In-text references to the RBDs omit the country code prefix (BE = Belgium) and the region code suffix (RW = Walloon Region, VL = Flanders).

**Table 2.** Overview of the pesticides used in this study, with the respective substance group (after Porta et al., 2025). HC5 = hazardous concentration to 5% of the species.

| Substance no. | Substance name | Substance code | Substance group | CAS Code | HC5 (µg/L) |
| --- | --- | --- | --- | --- | --- |
| 1 | Chlorothalonil | chloro | Fungicide | 1897-45-6 | 0.03650 |
| 2 | Pyraclostrobin | pyracl | Fungicide | 175013-18-0 | 0.3409 |
| 3 | Tebuconazole | tebuco | Fungicide | 107534-96-3 | 9.562 |
| 4 | Thiophanate-methyl | thioph | Fungicide | 23564-05-8 | 65.13 |
| 5 | Trifloxystrobin | triflo | Fungicide | 141517-21-7 | 1.319 |
| 6 | 2,4-Dichlorophenoxyacetic acid | 24dich | Herbicide | 94-75-7 | 86.50 |
| 7 | Dicamba | dicamb | Herbicide | 1918-00-9 | 103.8 |
| 8 | Dichlorprop | dichlo | Herbicide | 120-36-5 | 45.03 |
| 9 | Dimethenamid(-p) | dimete | Herbicide | 163515-14-8 | 92.85 |
| 10 | Diuron | diuron | Herbicide | 330-54-1 | 0.3887 |
| 11 | Glyphosate | glypho | Herbicide | 1071-83-6 | 86.14 |
| 12 | MCPA | mcpaaa | Herbicide | 94-74-6 | 314.9 |
| 13 | Pendimethalin | pendim | Herbicide | 40487-42-1 | 0.2128 |
| 14 | Chlorpyrifos | chlorp | Insecticide | 2921-88-2 | 0.008966 |
| 15 | Dimethoate | dimeto | Insecticide | 60-51-5 | 20.00 |

The overall objective of this study was to develop and evaluate a multi-level Bayesian network applicable for probabilistic risk characterisation of chemical mixtures in European river basins. The specific aims of this paper are: (1) to introduce and explore a novel probabilistic approach to mixture risk characterisation, which combines the CA and IA concepts in a multi-level BN; (2) to assess the ability of the BN to rank substances and substance groups according to risk contribution. Based on these evaluations, we consider the usefulness of this approach beyond the pilot study: for expansion to larger geographic scales, higher spatial resolution, higher number of substances, and other scenarios.

## 2. Modelling approach

### 2.1. Pilot study

As a pilot study for developing the Bayesian network modelling approach, we have selected a set of pesticides used in Belgium for the year 2012. The spatial and temporal settings of the current BN (referred to as Pilot 3) corresponds to those of a previously published BN (referred to as Pilot 2) (Mentzel et al., 2026). The Pilot 2 study described the overall probabilistic framework of the ENCORE project, as well as Bayesian updating of predicted concentrations for individual pesticides. In comparison, the current paper focuses on the modelling of mixture risk, with a revised selection of pesticides compared to Pilot 2. (Pilot 1 represents initial testing based on PECs from the project SOLUTIONS (van Gils et al., 2020), and will not be further described here).

The starting point for this BN is the chemical concentrations predicted by the ENCORE fate model: a continental-scale process-based simulation system based on the modelling framework of SOLUTIONS (van Gils et al., 2020), and further developed in the ENCORE project (Vlaeminck et al., 2026). This model system will in the following be referred to as the exposure model. It predicts environmental concentrations for a broad range of substances, including pesticides, pharmaceuticals, REACH chemicals, and legacy pollutants. The model integrates harmonised input data, advanced emission modelling, hydrological connectivity, and stochastic Monte Carlo simulation to account for uncertainty.

The spatio-temporal extent and resolution of the BN in Pilot 3 correspond to those of Pilot 2: the spatial units are the five river basin districts (RBDs, Figure 1), while the temporal units are the 12 months (Figure 2b). In contrast, the exposure model (Vlaeminck et al., 2026) has a higher resolution: the spatial resolution is ∼23,000 hydrologically connected subcatchments (144 for Belgium; Supplementary File 1), while the temporal unit is days. (Mentzel et al., 2026) provide more details on the spatial mapping of subcatchments to RBDs, and further reflections on the suitability of the BN model as an emulation model ENCORE exposure model (or fate model), here referred to as the exposure model.

**Figure 2.**
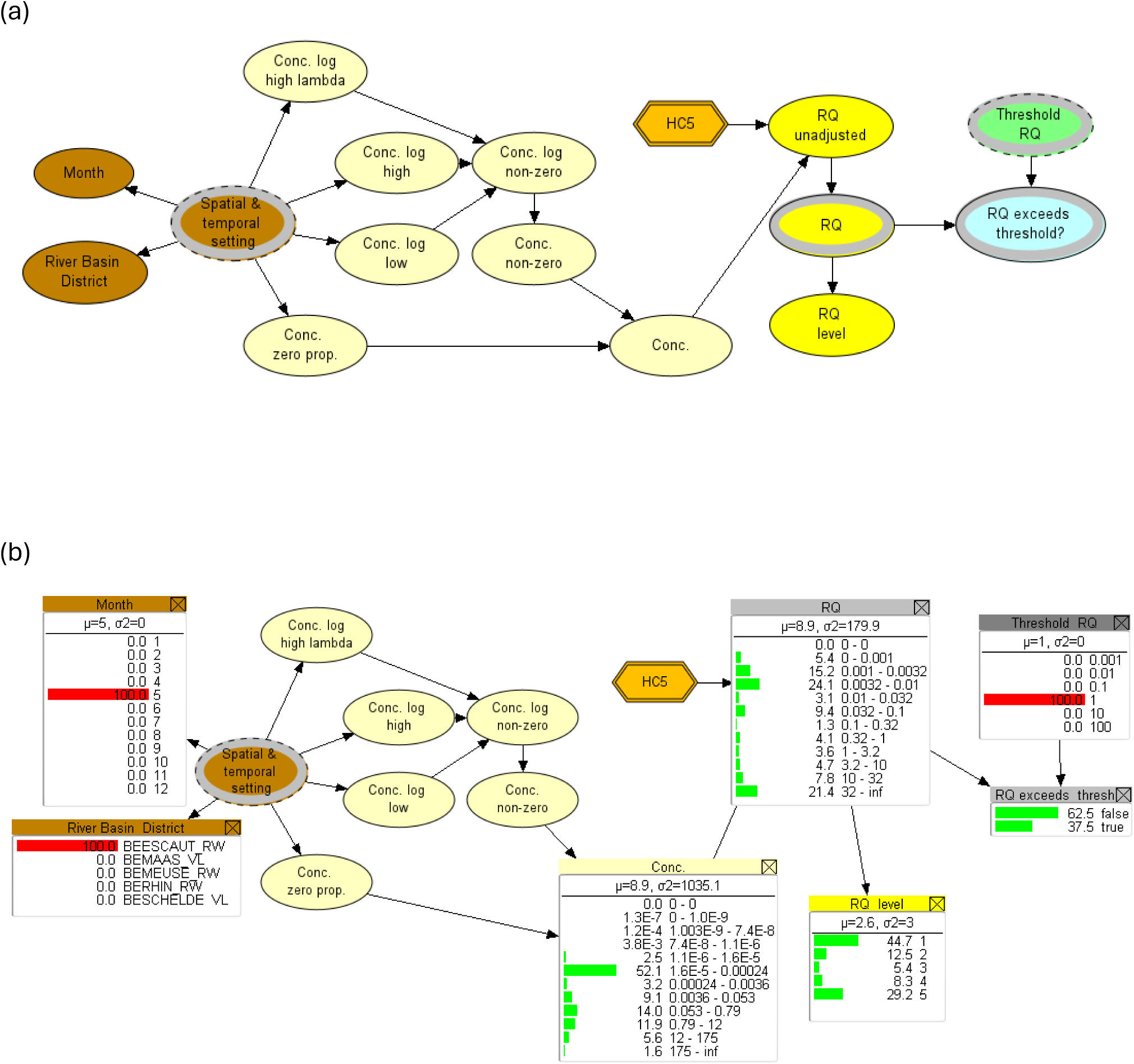
Submodel at level 1: Substance level. Example: “Subst_chloro”. (a): Graphical structure of the submodel (class) in the BN software. (b): Example of settings for scenario nodes (“River Basin District” = BEESCAUT_RW, “Month” = 5 (May), “Threshold RQ” = 1) and resulting probability distributions for a selection of nodes (“Conc.”, “RQ”, “RQ level”, and RQ_exceeds_threshold”. The monitor windows display states to the right (intervals, categories, etc.)., and the probabilities to the left (as horizontal bars and percentages). For a description of each nodes, see Table 4a. Nodes with thick grey outlines represent connections to other submodels: dashed lines show input nodes (e.g. “Threshold RQ”), solid lines show output nodes (e.g. “RQ”).

**Table 3.**
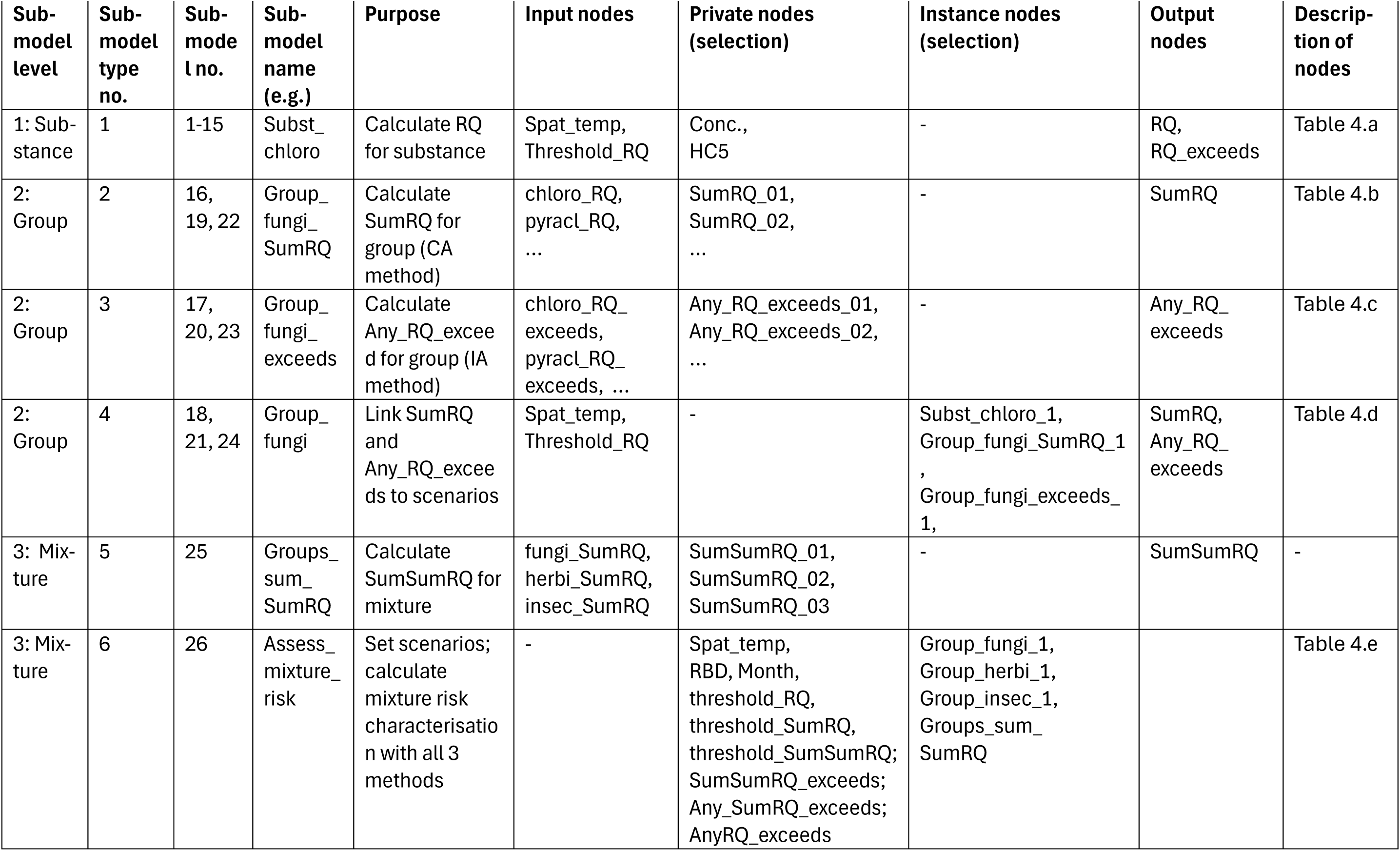
Overview of submodels in the object-oriented Bayesian network. For more information in the individual nodes, see Table 4.

**Table 4.**
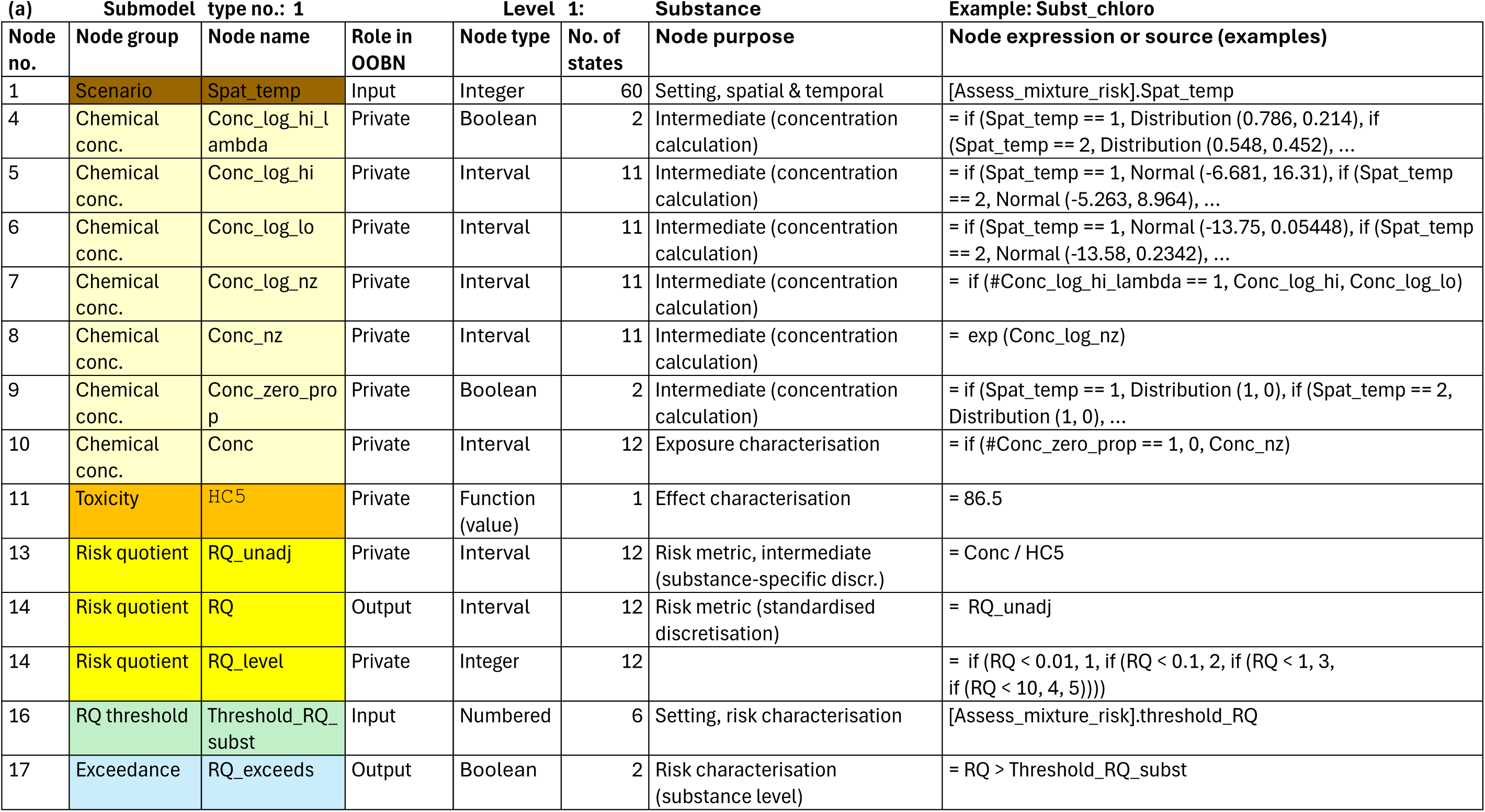
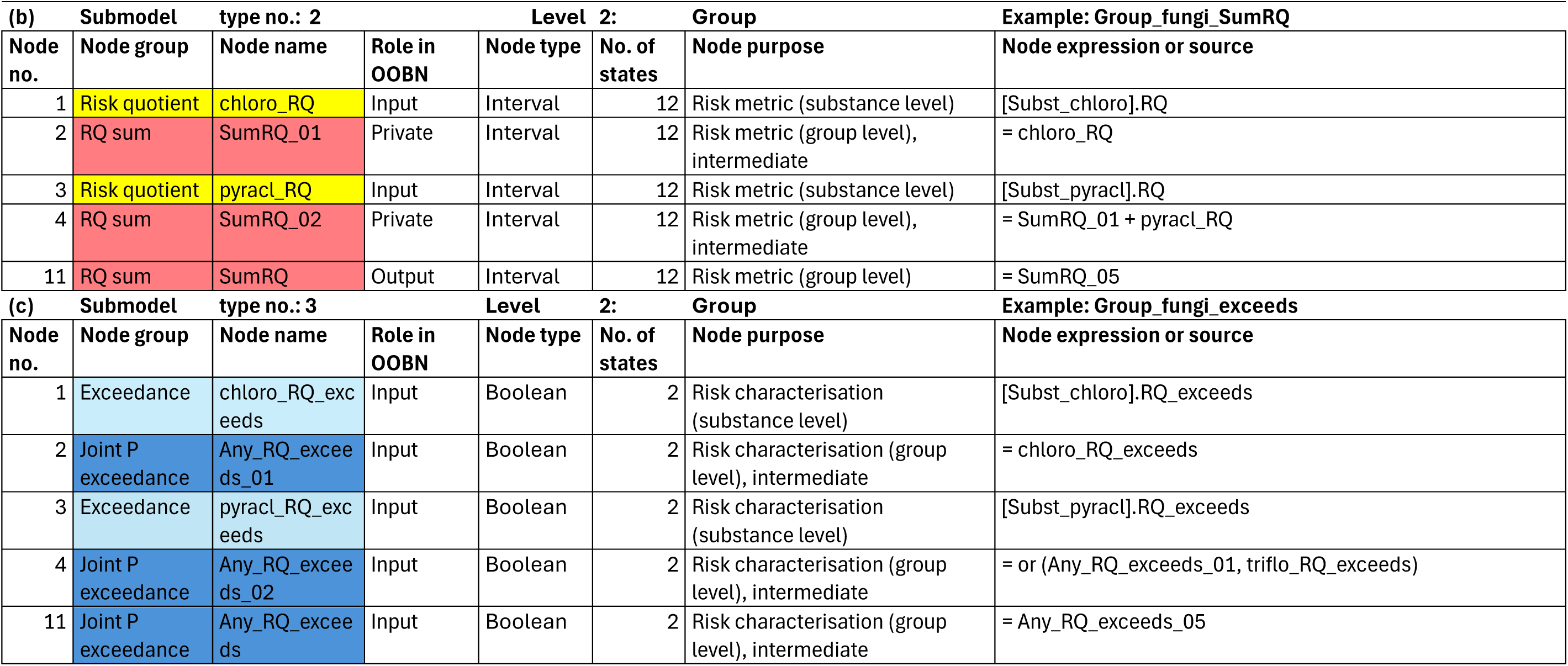
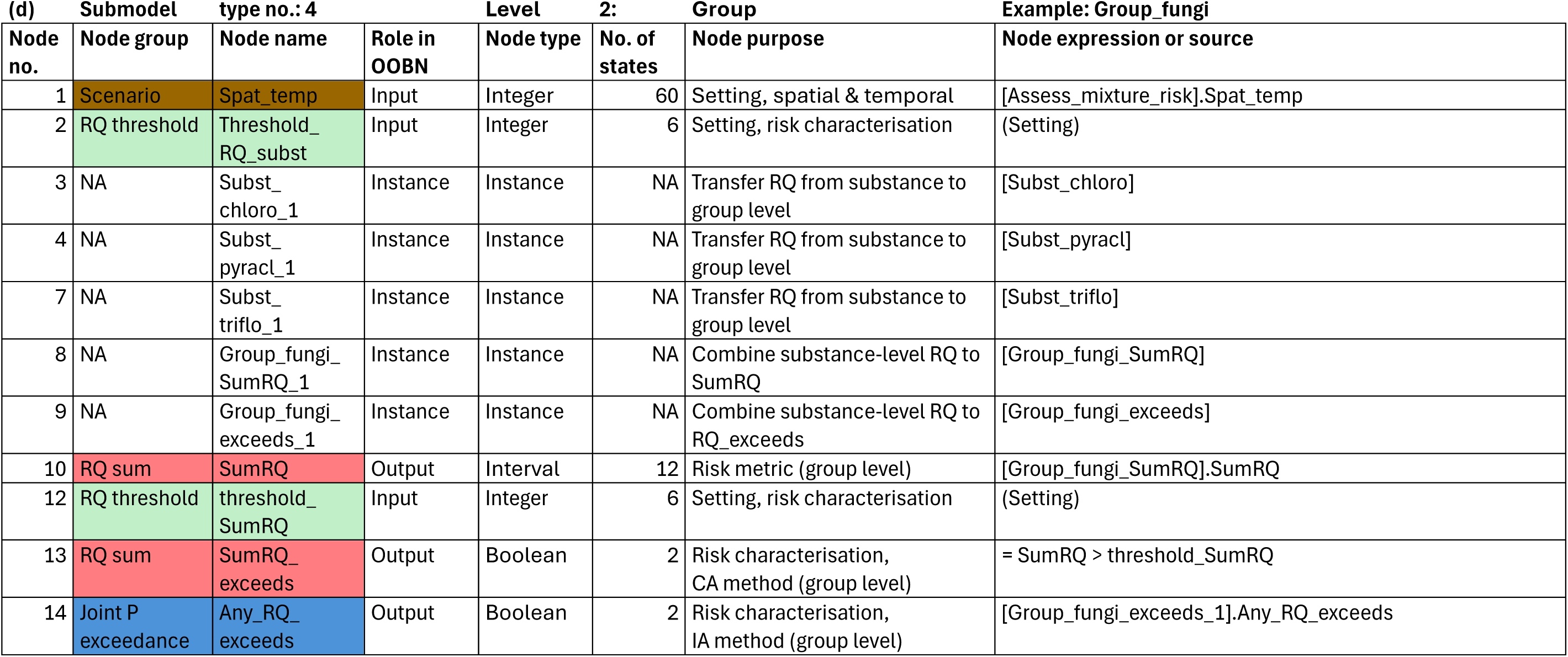
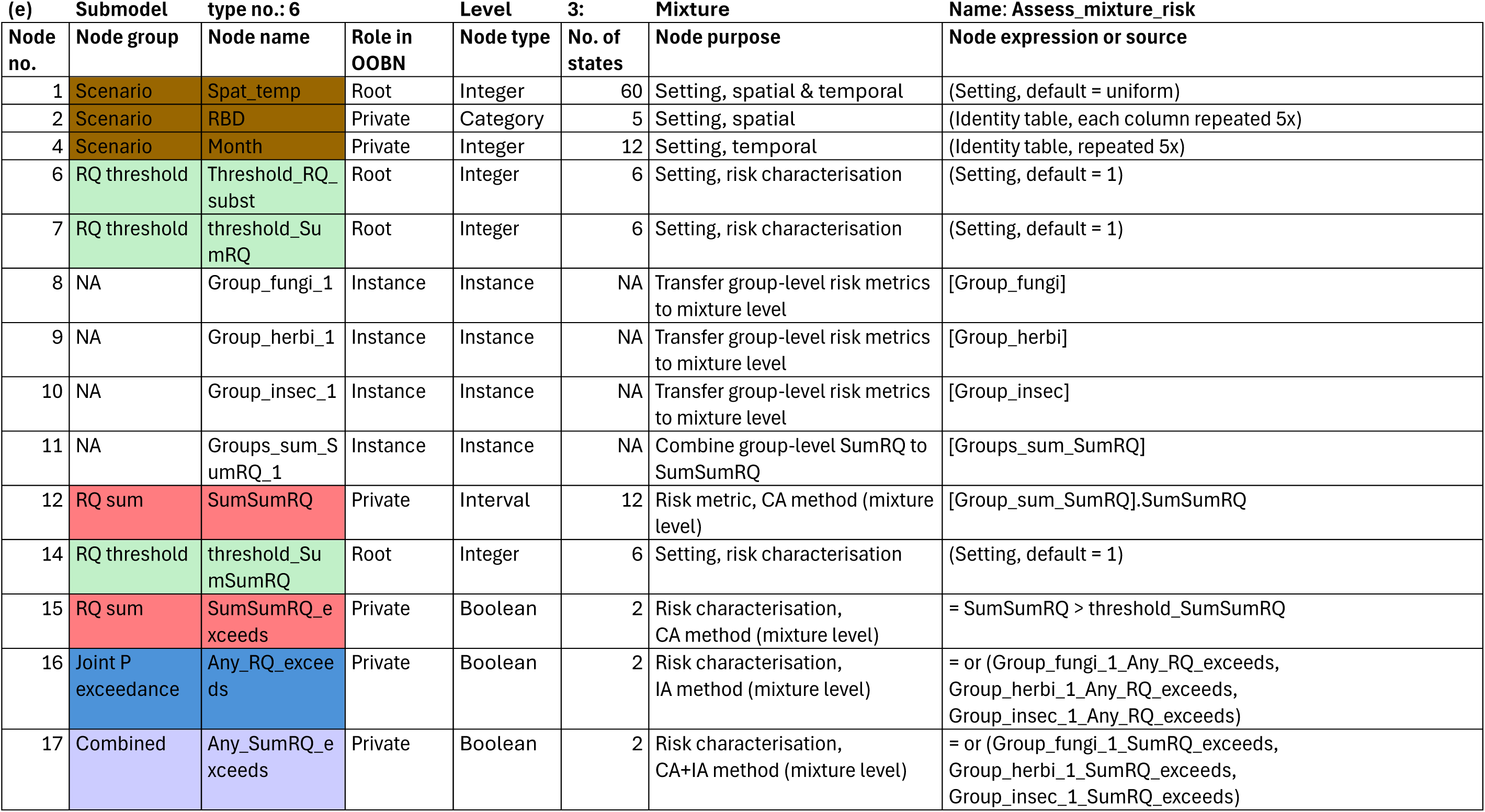
Overview of all nodes in the 3-level Bayesian network. The list of nodes is organised according to the submodels in Table 3. Level 1 Substance: (a) Submodel type no. 1, example “Subst_chloro” (Figure 2). Level 2 Group: (b) Submodel type no. 2, example “Group_fungi_SumRQ” (Figure 3a); (c) Submodel type no. 3, example “Group_fungi_exceeds” (Figure 3b); (d) Submodel type no. 4, example “Group_fungi ” (Figure 3c). Level 3 Mixture: (e) Submodel type no. 6, “Assess_mixture_risk” (Figure 4).

The selection of 15 pesticides in Pilot 3 (Table 2) differs from the selection in Pilot 2, and was primarily based on the availability of monitoring data in Waterbase (Agency, 2025). The calculated use of each pesticide, according to the methodology of Porta et al. (2025), is the average use (in kg/year) over the years 2011 to 2017, using national sales data from Fytoweb as proxy. As described for Pilot 2 (Mentzel et al., 2026), the main input for the BN is the PECs obtained from the ENCORE exposure model (Vlaeminck et al., 2026). The exposure model version used for the current study is a refined version compared to Pilot 2: the main difference is the inclusion of newer high-resolution application rate maps for Europe (Porta et al., 2025). The further processing of the PECs used in the current study (Supplementary File 2) is described in Section 3.1.3.

### 2.2. Mixture risk characterisation: concepts and probabilistic methods

The single-substance risk characterisation in this study is based on the concept of Risk Quotients (RQ), which represents the ratio of an environmental concentration to an environmental threshold value (ETV). The environmental concentration can be predicted (PEC), as in this study, or measured (MEC) (or a combination, as exemplified by Mentzel et al. (2026). RQs and similar metrics are often reported as single values or by basic summary statistics for a given area. Here, in contrast, RQs are quantified by full probability distributions for any spatio-temporal unit.

The environmental threshold value for a given substance is often a PNEC (predicted no-effect value), which can be derived in various ways, but is often calculated as the HC5 value (hazardous concentration for 5% of the species) divided by an assessment factor to account for uncertainty by lowering the threshold. In ENCORE, we use the HC5 value directly as the ETV without additional protective assessment factors, since the probabilistic framework allows for more transparent handling and propagation of uncertainty. The HC5 values (Table 2) were obtained from (Posthuma et al., 2019), as for Pilot 2. RQs have been widely used for screening-level risk assessments of individual substances by checking whether environmental thresholds are exceeded, i.e. whether the RQ (= PEC/ETV) exceeds a value of 1. In ENCORE, we have expanded this procedure to check whether RQ exceeds a range of threshold levels (Threshold_RQ, ranging from 0.001 to 10), while still referring to the condition RQ > 1 as the default threshold exceedance.

The mixture risk calculation in this study is also based on RQ calculation. However, while other examples of RQ-based mixture risk calculation normally represent the CA approach (Table 1), our study makes use of RQs for all three variants of mixture risk calculation (CA, IA and CA+IA). CA is a component-based model that assumes that all components have a similar mode of action (MoA), e.g. by affecting the same molecular targets in an organism, even if they differ in toxic potency. Each component contributes to the overall effect based on the ratio of its concentration to its toxic potency (EFSA Scientific Committee et al., 2019).

In ecotoxicology, IA (independent action) is a component-based model that assumes that all components are independent (dissimilar action), following the statistical concept of independent random events. (The concept has also been referred to by alternative terms such as “response addition”) (de Zwart and Posthuma, 2005) and “effect addition” (Liess et al., 2016)). The traditional application of IA in ecotoxicology requires dose-response data at organism level (e.g. mortality) to be expressed as a proportion (between 0 and 1). This number can represent, for example, the proportion of individuals in a test population being killed by the mixture exposure (EFSA Scientific Committee et al., 2019). Here, we apply the calculation used with the IA concept in a more general sense, to represent joint probabilities of independent events. The “action” or “event” in this context is the exceedance, RQ > Threshold_RQ. With this interpretation of the IA concept, we assume that chemicals assigned to different chemical groups pose risks to ecosystems in an independent manner. Therefore, the combined risk can be calculated by cumulative probability of effect threshold exceedance, rather than by cumulative toxic effects on specific species or target organs.

In this study, thus, risk characterisation for mixtures was calculated using all of the three methods (Table 1): (1) CA, the sum of risk quotients for all substances (the node “SumRQ_exceeds”), (2) the combined CA+IA method (“Any_SumRQ_exceeds”), and (3) IA, the joint probability of RQ threshold exceedance for one or more substances (“Any_RQ_exceeds”).

The combined method (CA+IA) represents a three-step calculation, where the last two steps correspond to the two-stage prediction (TSP) model of Junghans (2004) and collaborators. First, probabilistic risk quotients (RQ) are calculated for each individual substances, as a probability distribution of environmental concentrations (RQ = PEC/HC5). Next, the CA concept is applied within groups of substances (presumed to have similar modes of action), by summing the RQ distributions (SumRQ). Finally, the IA concept is applied to SumRQ distributions across the different substance groups, by calculating the joint probability of threshold exceedance (SumRQ > Threshold_RQ) for one or more groups.

### 2.3. Bayesian network methodology

A Bayesian network (BN) is a graphical probabilistic model that represents a set of variables and their conditional dependencies via a directed acyclic graph. The edges (links) can have a causal interpretation and a corresponding set of conditional probability distributions, which factorises the joint probability distribution over the variables of the model (Kjærulff and Madsen, 2013). The variables, referred to as nodes, are usually discretised into intervals or categorical states, although hybrid BNs can also contain continuous nodes (Moe et al., 2020). The arrows pointing from one or more parent nodes to a child node represent a relationship, preferably with a causal structure, which is quantified with a conditional probability table (CPT). The CPT relates the probability for each state of the child node to each of the states of the parent nodes. The BN uses the CPT of a child node in combination with the probability distributions of its parent nodes to calculate the probability distribution of the child node. This calculation is done according to Bayes’ rule, which describes the probability of an event conditional on prior knowledge of conditions that might be related to the event (Kjærulff and Madsen, 2013). In Bayesian inference, thus, the posterior probability (prediction) for a node is proportional to the product of the prior probability (assumptions) and the likelihood function (new evidence). A well-known advantage of this approach is that BNs can be constructed from tentative information sources (prior probability distributions) and updated/refined with independent data (e.g., monitoring data) (e.g., Mentzel et al., 2026). In this paper, however, we focus on the use of BNs as a tool for probabilistic mixture risk calculation, applied to both CA-based and IA-based methods as well as the combination CA+IA (Table 1).

An important property of BNs, compared to other methodologies for probabilistic modelling, is that a BN model can have both deterministic and probabilistic properties. For example, if a BN model is run 100 times with identical settings, the outcome for a given variable (e.g. chemical concentration) will be 100 identical probability distributions. In comparison, for a process-based model with a stochastic component (e.g. Monte Carlo simulation), the outcome of 100 runs for a given concentration variable will be 100 (slightly) different values, which can be expressed as a single probability distribution. While the resulting probability distribution can be similar in the two situations, there is a fundamental difference between the two approaches: the deterministic property of the BN allows for additional types of diagnostic inference, such as sensitivity analysis, directly from the parameterised model by exact probability calculations (e.g., Moe et al., 2020). In comparison, sensitivity analysis for a stochastic process-based model would typically require simulations with multiple model runs. More details on sensitivity analysis for the BN developed here is given in Section 3.5. Another advantage of BN methodology is the opportunity for combining submodels (technically term: “classes”) into multi-level networks (object-oriented BN) Table 3. More details on the OOBN structure and development for this study are given in Section 3.2.

The starting point for mixture risk calculation with the BN in this study is the RQ, expressed as a discrete probability distribution across 12 intervals (Table 4a), for each of the 15 substances. The CA method can be implemented in a BN simply by adding the RQ distributions (Eq.1):

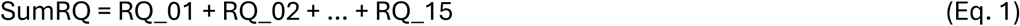

While the equation is straight-forward, the resulting SumRQ can to some degree be affected by discretisation of each RQ node (the handling of this issue is described in Section 3.3).

The IA method, which represents the joint probability of multiple independent events, has been applied in ecotoxicology to express the probability of a certain effect (e.g., exceeding 50% mortality) for a given combination of substances (*i*) each with concentration *c_i_*. The equation can be generalised for a mixture of *n* substances (Eq. 2, after Backhaus et al. (2004)):

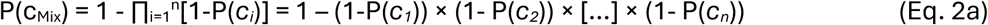

While the IA method was initially formulated for binary response data (e.g. mortality vs. survival), it has been generalised to apply for continuous response variables, with effect (*E*) expressed as a proportion (range 0 -1) or percentage (0-100%) (Eq. 2b, after Backhaus et al. (2004)):

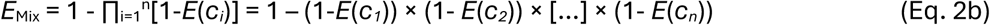

Eq. 2b (or 2a) can be implemented in a BN effectively by a Boolean “*OR*” expression, stating the joint probability that one or more of the events *E_i_* will occur (Eq. 3a):

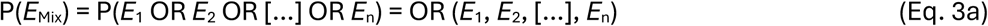

For the current study, the individual event ***E_i_*** is defined as the situation **RQ*_i_* > Threshold_RQ** (labelled “RQ_exceeds”). The joint probability of RQ exceeding the threshold for one or more substances (labelled “Any_RQ_exceeds”) can then be expressed as Eq. 3b:

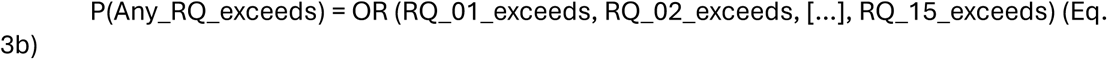

When implemented in a BN software, this expression will be translated to conditional probability table (CPT) for the target node (Any_RQ_exceeds) (see Supplementary File 6).

More specific information on the implementation in of CA and IA as well as the combined CA+IA in the multi-level BN developed here, are given in Section 3.4.

## 3. Development of a Bayesian network for mixture risk calculation

### 3.1. Workflow and data processing

#### 3.1.1. Pilot BE pesticides

The principles and steps of the workflow for this pilot study largely follow the description of Pilot 2 (Mentzel et al., 2026). The main novelty of Pilot 3 compared to Pilot 2 (as explained in Section 2.1) is the combination of probabilistic risk characterisations (RQ distributions) from the single substance level into the group and mixture levels. Moreover, a procedure for refined discretisation of concentration and RQ nodes has been developed for Pilot 3. The current Section will therefore focus on the steps involved in constructing the multi-level BN for mixture risk characterisation, as well as the steps for refining the submodels for single-substance risk characterisation.

#### 3.1.2. Work flow

The main input data are the PEC values simulated by the ENCORE exposure model (described in Section 2.1), available as a set of 15 text files (comma-separated values - csv, Supplementary File 2). Each file contains the 526,032 simulated concentration with unit g/m^3^ (= 0.001 µg/L), representing 144 subcatchments x 365 days x 10 years (2004-2013) for one pesticide. Initial processing of the PEC values, including compilation, conversion to unit µg/L, combination with HC5 values and RQ calculation, were carried out in Microsoft Access for initial data screening. All further data processing and statistical analyses to support the BN development were carried out in the programming environment R version 4.5.2 (R Core Team, 2025).

The BN model was constructed, run and analysed in the software HUGIN Researcher version 9.7 (Madsen et al., 2005). The complete model code is provided as a text file (Supplementary File 5), which can be opened with the HUGIN software. The software-generated model documentation provides more user-friendly information and illustrations in interactive html files (Supplementary File 6). An publicly available online user interface to the BN (https://encore.hugin.com/models/Pilot) has been developed to allow any users to explore the main scenarios and outcomes, without installing the HUGIN software (see Supplementary File 7).

#### 3.1.3. Statistical distributions

For aggregation of the simulated PEC values from the original spatio-temporal resolution (subcatchment and day) to the BN’s resolution (RBD and month), statistical distributions were fitted to the PECs for each RBD and month. The purpose was to obtain a proper statistical representation of the variability of PECs at this scale, instead of collapsing the information to simpler summary statistics such as a mean or median, as recommended by Moe et al. (2022). Previous studies involving probabilistic exposure characterisation have often used lognormal distributions, which can be a reasonable approximation for single-substance risk characterisation (e.g., Mentzel et al., 2022). Since the lognormal distributions is strictly positive, zero concentrations (and consequently zero RQ) will typically be approximated with very small positive number; this is normally not a problem when considering the risk of a single substance. However, when RQ values are combined for mixture risk assessment, small deviations from zero can accumulate and result in a spurious over-estimation of mixture risk. This is especially the case when applying the CA method with discretised RQ values (see Tollefsen et al., 2025). Moreover, visual inspection of histograms for the PECs (Supplementary File 3) revealed conspicuous bimodal distributions, which would require more sophisticated distributions than lognormal. The bimodal patterns were consistent for most of the different pesticides, RBDs and months, although the amplitudes varied throughout the year. These patterns could be explained by various settings in the process-based model (Vlaeminck et al., 2026), with the dominant factors being a combination of seasonal pesticide applications and hydrological and chemical processes (transport, degradation etc.).

The solution to fitting the bimodal patterns in PEC values was a four-step procedure. First, concentration values below a given limit (10^-9^ µg/L) were rounded to zero. Second, the concentrations were split into zeros and positive values. Third, the positive concentrations values were log-transformed (ln, natural logarithm). Fourth, a statistical mixture model representing normal distribution with two components was fitted to the log-transformed concentration, using the R package “mixtools” (Benaglia et al., 2009). (The term “mixture” here refers to the distribution as a mixture of two statistical distribution components, with no relation to chemical mixtures). Although lognormal distributions could have been fitted directly to the un-transformed values, we chose an additional step of log-transformation before fitting normal mixture distributions, for transparency. The purpose was to obtain a stepwise processing that could be easily implemented in the BN (Section 3.4.2). The resulting plots (Supplementary File 3) showed that the two components of the mixture model in general corresponded well to the two peaks of the PEC histograms. The fitted parameters are listed in Supplementary File 4: lambda (weight of each component), µ (mean) and sd (standard deviation), for each of the 900 distributions (15 substances x 5 RBDs x 12 months).

The 15 pesticides were assigned to three different substance groups, according to their chemical nature and functional mode of action (based on Porta et al., 2025): fungicides (including bactericides), herbicides (including haulm destructors and moss killers), and insecticides (including acaricides). This grouping reflects differences in application practices and intended targets, which act on specific components and developmental stages within the ecosystem. We have assumed that the environmental risk in different substance groups, i.e. the exceedance of RQ thresholds, can be considered independent events across the different groups. In principle, this implies that the concentration of different substance groups should be resulting from independent processes (chemical use and emission), and that the toxic effect of different substance groups should be independent (i.e., different mode of action). While the toxic mode of action concept is originally meant for effects on species and species groups (EFSA Scientific Committee et al., 2021), it is here applied in a more general sense to derived risk metrics such as RQ. Thus, for simplicity and for the sake of method demonstration, we have assumed that environmental risk (i.e., exceedance of RQ thresholds) in different substance groups can be considered as independent events in this study.

### 3.2. BN construction: Submodels, nodes and relationships

A multi-level or object-oriented Bayesian network model was developed for the pilot study. The full model code is available in the Supplementary Material (File 5). The BN has general structure of three main hierarchical levels: 1) Substances, 2) Groups, 3) Mixtures (Table 3), each consisting of multiple subnetworks, which will be referred to as submodels (technical term: “classes”). The level of a submodel is defined according to the use of output nodes from lower levels: Output nodes from submodels in Level 1 (Substance) are used as input nodes in Level 2 (Group); output nodes from Level 2 are used as input nodes in Level 3 (Mixture). The term «Mixture» is here used to mean all substances selected to be included in the BN model for the given pilot study (as opposed to the subset of substances assigned to a specific substance group). For each of the three levels, a selection of representative submodels are explained (Table 4) and illustrated (Figure 2, Figure 3a,b,c, Figure 4). For the same selection of submodels, system-generated model documentation is provided as a set of html files (File 6).

**Figure 3.**
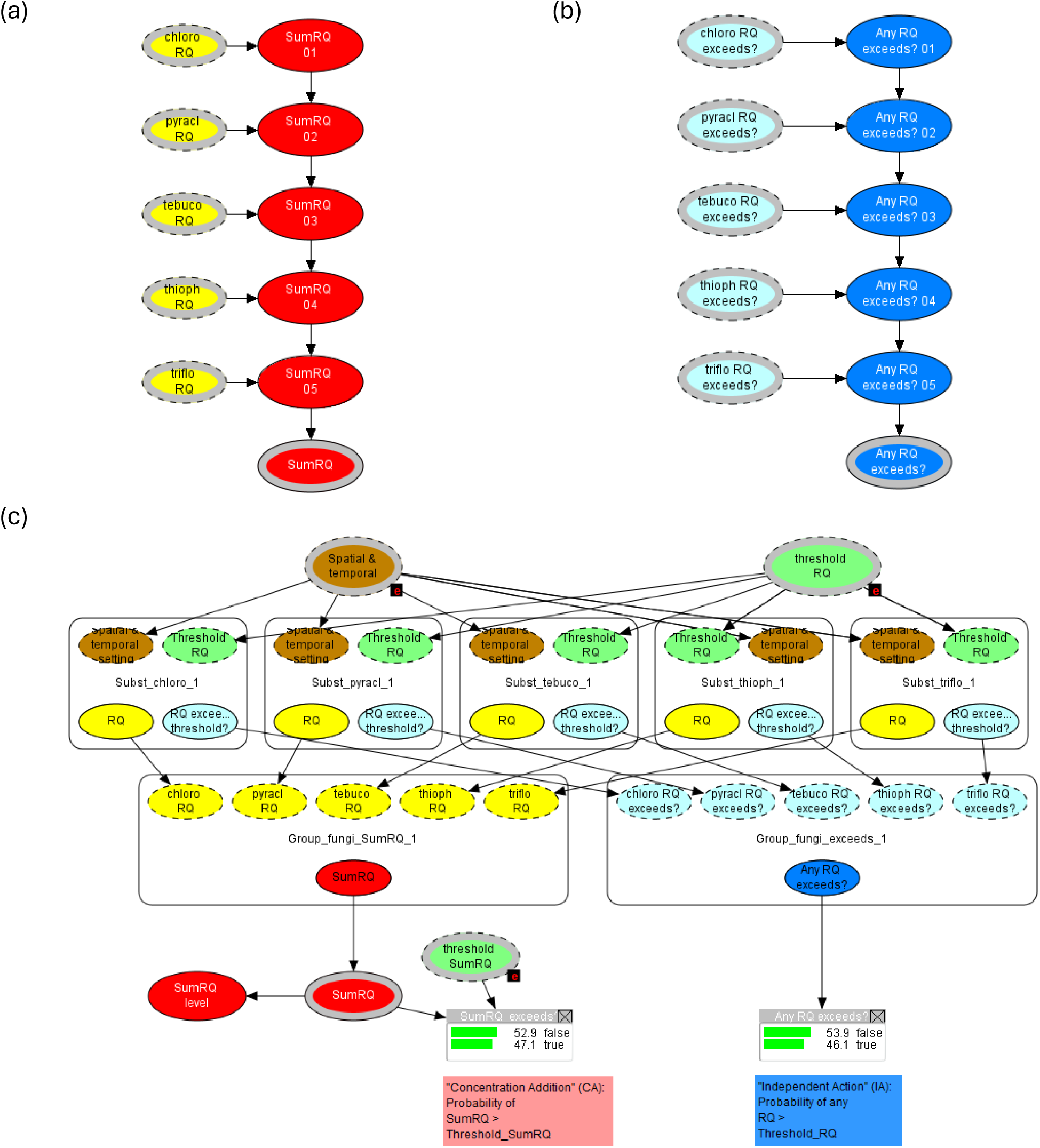
Submodels at level 2: Group level. Examples shown for the substance group “fungi” (Fungicides). a) Submodel “Group_fungi_SumRQ”, for CA-type risk calculation (Submodel type 2, see Table 4b). (b) Submodel “Group_fungi_exceeds”, for IA-type risk calculation (Submodel type 3, see Table 4c). (c) Submodel “Group_fungi ” (Submodel type 4, see Table 4d). Here, the relevant submodels for fungicide substances are included as instance nodes. For example, the instance node “Subst_chloro_1” represents an instance of the submodel Subst_chloro (Figure 2). The instance nodes “Group_fungi_SumRQ_1” and “Group_fungi_exceeds_1” represent submodels for group-level risk calculation by the CA and IA concepts respectively (see Table 1). Probabilities are shown for the output nodes “SumRQ exceeds?” (for group-level CA) and “Any RQ exceeds?” (for group-level IA). Black boxes with the label “e” indicates that evidence has been set for the node (corresponding to the red bars in Figure 2b. (See Figure 2 for more information on input and output nodes).

**Figure 4.**
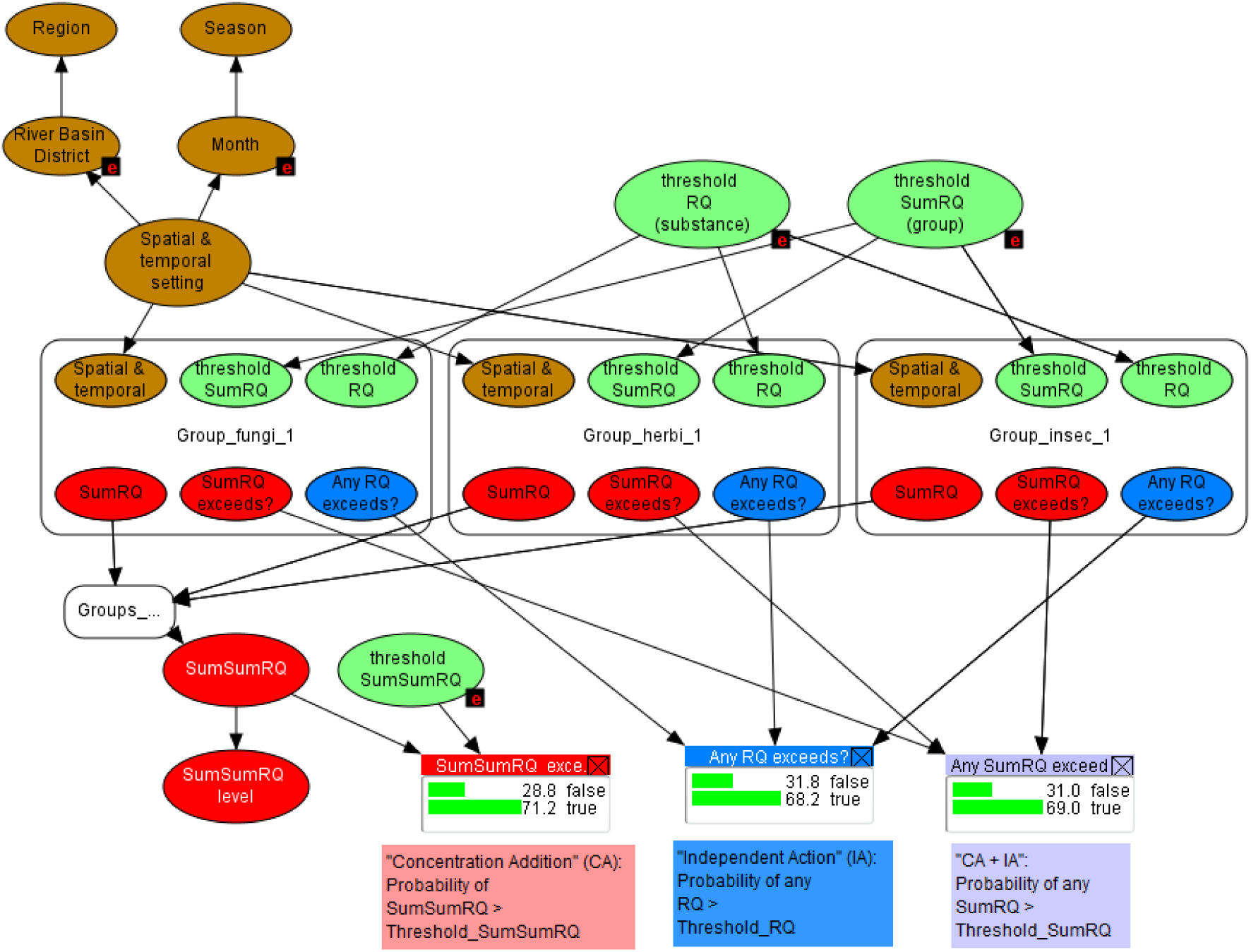
Level 3 sub-model: Mixture level (top level). Submodels from the group level representing all three groups are included as instance nodes. For example, the instance node “Group_fungi_1” represents an instance of the submodel Group_fungi (Figure 3). The collapsed instance node “Groups_sum_SumRQ_1” serves as a sub-model to simplify addition of SumRQ from all groups. See Figure 2 for input node settings, and for explanations to input and output nodes. For explanation of the three methods for mixture risk calculation, see Table 1).

The submodels are connected in higher level submodels by input and output nodes, which are indicated by thick grey outlines in combination with dashed black outlines (input) or whole black outlines (output) (e.g. Figure 2A). To facilitate the visual interpretation, all nodes are assigned to a set of thematic node groups with different colours, as described in Table 4 and displayed in illustrations of the BN (Figure 2–Figure 4).

More information on states, discretisations, prior probability distributions and expressions are given in the subsequent tables. The main nodes and node types are described in Table 3, while more complete node descriptions are given in (Table 4) and in Supplementary Material File 7. The R package RHugin (Konis, 2026) was used as a middleware between R and HUGIN for efficient creating of submodels for all substances, which had identical structure but different parametrisation.

#### 3.2.1. Level 1: Substances

Level 1 has one type of submodel (named “**Subst_xxxxxx**”, Table 3a), used for each of the 15 substances (Table 2), and exemplified for the substance Chlorothalonil (Subst_chloro). For all substances the subnetwork has identical structure, but different discretisation and quantification for some of the nodes. The main purpose of this node is to quantify the risk quotient (RQ) by a probability distribution.

##### Node group Scenario

The substance subnetwork contains the input node labelled “Spatial & temporal setting” (node name: Spat_temp), with numbered states 1-60. This node combines all possible states of the two scenario nodes, the spatial scenario node labelled “River Basin District” (node name: RBD; 5 states; see Figure 1) and the temporal scenario node “Month” (12 states). The input role of Spat_temp means that this node can be instantiated from another submodel (described in Section 3.2.3).

##### Node group Chemical concentration

The chemical exposure concentration (node “Conc.”) is calculated from a series of nodes representing different components of the estimated statistical distribution; these procedures are described in Section 3.4.

##### Node group Toxicity threshold

The node HC5 is a single-value function node, containing the substance-specific toxic effect metric as listed in Table 2. The HC5 value is the estimated hazardous concentration to 5% of species, obtained from Van Gils et al. (202x).

##### Node group Risk Quotient

The risk quotient (RQ) is calculated as the exposure concentration divided by the effect threshold (here: Conc/HC5). The two node variants RQ_unadjusted and RQ differ only by the discretisation: the internal node RQ_unadjusted has discretisation specific for each substance (determined by the range of Conc), while the output node RQ has discretisation adjusted to a common scale across all substances (see Section 3.3 for more details). The node RQ is an output node, used for multi-substance risk calculation by the Concentration Addition method at the next level (see Table 1). The third variant, “RQ level” (5 numbered states), is used for extracting model predictions, and have no further role in the BN.

##### Node group RQ threshold

The node «Threshold RQ» is a numbered node with six alternative RQ threshold values (ranging from 10^-3^ to 10^2^), which can be selected for risk calculation. The default value is 1, representing the situation where the exposure concentration exceeds the environmental threshold value.

##### Node group Exceedance

The Boolean (false/true) node labelled “RQ exceeds threshold?” represents the probability of RQ exceeding the selected RQ threshold. This output node is used for multi-substance risk calculation by the Independent Action method at the next level.

#### 3.2.2. Level 2: Groups of substances

Level 2 contains three types of submodels (Table 3b-d), each used for each of the three substance groups. The first two represent the CA and IA calculation methods, respectively, while the third makes use of these as instance nodes.

##### Submodel type 2: Group_xxxxx_SumRQ

(Table 3b), exemplified by Group_fungi_SumRQ (Figure 3a). This submodel is used for calculating the sum of RQ for all substances in a group by sequential adding of RQ nodes. The input nodes (chloro_RQ, pyracl_RQ, etc.) are RQ distributions for different substances. The internal nodes (SumRQ_01, SumRQ_02, etc) contain the cumulative sum of these inputs. Lastly, the final output node SumRQ makes this risk metric available for other submodels that make used of the CA method (Figure 3c). The nodes are organised sequentially instead of in parallel, in order to limit the size of the conditional probability tables (Section 3.4): for each additional parent node, the number of columns in the current CPT must be multiplied with the number of states of the new parent (here: 12).

##### Submodel type 3: Group_xxxxx_Any_RQ

(Table 3c), exemplified by Group_fungi_Any_RQ (Figure 3b). This submodel is used for calculating the joint probability of RQ exceeding the threshold for multiple substances (see Table 1), from the Boolean input nodes (“chloro_RQ_exceeds”, “pyracl_RQ_exceeds”, etc.). As for Group_fungi_SumRQ, the input nodes are organised sequentially to limit the size of the CPTs, although the parent nodes now have only 2 states. The resulting output node, “Any RQ exceeds”, represents the joint probability that RQ exceeds the given threshold for any of the substances involved in the calculation. (More details on the joint probability calculation are given in Section 3.4).

##### Submodel type 4: Group_xxxxx

(Table 3d), exemplified by Group_fungi (Figure 3c). The purpose of this submodel is to connect the submodels for group-level calculations (Group_fungi_SumRQ, Group_fungi_Any_RQ) to the individual substances, as well as to settings and to further risk calculations. This is obtained by instance nodes (displayed as rounded rectangles with nodes inside), which represent instances of other subnetworks. The submodel Group_fungi contains instances of submodels for the five substances belonging to this substance group (labelled Subst_chloro_1, Subst_pyracl_1, etc), as well as the two submodels for group-level risk calculation (Group_fungi_SumRQ_1 and Group_fungi_Any_RQ_1). Additionally, this submodel contains nodes for calculating the probability that the calculated SumRQ exceeds the given threshold (node Threshold_SumRQ).

#### 3.2.3. Level 3: Mixtures of substances

##### Submodel type 5: Groups_sum_SumRQ

The purpose of this submodel is sequential additional of the SumRQ from the three groups into a mixture-level SumSumRQ (Concentration Addition method), in the corresponding way as done from substance to group level by the submodel Group_fungi_SumRQ (Figure 3a, Table 3b). This way, the number of parent states configurations (CPT columns) per calculation is reduced from 12^3 = 1728 to 12^2 = 144. Because this submodel is analogous to Submodel type 2, it is not described in further detail. Considering the Independent Action method, a corresponding submodel has not been constructed for sequential calculation joint probability of exceedance for group to mixture level. The reason that organising the parent nodes sequentially in this case would gain little computational efficiency, when the number of parent states configurations in this case is only 2^3 = 8.

##### Submodel type 6: Assess_mixture_risk

The final top-level submodel (Figure 4, Table 3e), combines all the group-level risk metrics to into mixture-level risk characterisations, and links them to the scenarios and other settings (RQ thresholds). This apex submodel has four instance nodes representing the three versions of Submodel type 4 (Group_fungi_1, Group_herbi_1 and Group_insec_1), as well as Submodel type 5 (Groups_sum_SumRQ_1) as an intermediate step (displayed in collapsed format in Figure 4). The three final Boolean nodes SumSumRQ_exceeds, Any_SumRQ_Exceeds and Any_RQ_exceeds correspond to the three mixture risk calculation methods CA, IA and CA+IA (Table 1), with further details on the calculations given in Section 3.4.

### 3.3. Discretisation of continuous variables

The discretisation of continuous variables, which is an inherent property of BNs, can be a crucial step for BN model development (Marcot, 2017). The number of states should strike a balance between precision (resolution) and model size (no. of parent states configuration), and capture the main characteristics of the nodes’ distribution. For the current pilot study, certain aspects of the data and their intended use pose particular challenges for the discretisation. First, the PECs span very wide ranges, both across and within the three dimensions (spatial, temporal and chemical). Second, the PECs have complex distributions: fundamentally log-normal, but also displaying bimodal as well as zero-inflated patterns (described in Section 3.4). Discretisation with exponentially sized intervals (i.e., equidistant intervals in log-scale) could in principle handle the PEC patterns across the wide ranges. However, the CA method poses an additional challenge in this case: preliminary model explorations have shown summing of RQ nodes with exponential intervals can result in inflated RQ values (Tollefsen et al., 2025). In the current study, thus, a substance-specific discretisation procedure was developed to handle the full range of PECs (with buffers) as well as the zero values. The resulting discretisations are listed in Supplementary File 9.

For each substance, the discretisation was performed as follows. The main exposure concentration node “Conc.” has 12 intervals, including a proper zero interval [0 - 0] (to avoid spurious accumulation of low concentrations with the CA method). First, a lower limit was set (PEC_low = 10E-9), below which values were rounded down to zero (see Section 3.1.3). Second, the minimum and maximum values (PEC_min and PEC_max) were extracted from the non-zero values. Third, an upper limit was set for discretisation (PEC_upp), corresponding to 110% of PEC_max. Fourth, the PEC values were log_e_-transformed. The range from log(PEC_min) to log(PEC_upp) were divided into 8 equal steps. The discretisation of the non-zero concentration range in log_e_ scale (Node Conc_log_nz) is thus 11 intervals with the following boundaries: [- inf; log_e_(PEC_low); log_e_(PEC_min); log_e_(PEC_min) + 1×step; log_e_(PEC_min) + 2×step; …; log_e_(PEC_min) + 7×step; log_e_(PEC_upp); infinity].

The discretisation for node Conc (12 intervals) was obtained by back-transformation, and adding the first zero interval):

[0; 0; PEC_low; PEC_min; exp(log_e_(PEC_min) + 1×step); exp(log_e_(PEC_min) + 2×step); …; exp(log_e_(PEC_min) + 7×step); PEC_upp; infinity].

An example of discretisation for node Conc is shown in Figure 2b.

For the node RQ_unadjusted, the substance-specific intervals boundaries were derived from the Conc interval boundaries by dividing by HC5. For the node RQ, the interval boundaries were adjusted to be common for all substances, with equidistant steps in log_10_ scale:

[0; 0; 10E-3.0; 10E-2.5; …; 10E1.0; 10E1.5; inf] (also displayed Figure 2b).

The log_10_ scale was chosen for the RQ node to obtain a scale with straightforward interpretation. For concentrations, on the other hand, the log_e_ scale was chosen to facilitate the use of built-in expression components, such as log_e_-transformation.

The three variants of numbered nodes for RQ thresholds (“Threshold_RQ”, “Threshold_SumRQ”, “Threshold_SumSumRQ”) all had the same discretisation, namely a subset of the interval boundaries from the RQ nodes: [10E-3, 10E-2, 10E-1, 10E0 , 10E1, 10E2). The aggregated version RQ_level have numbered states 1-5, corresponding to the intervals between these 6 alternative RQ thresholds.

### 3.4. Prior distributions and conditional probabilities

The complete set of conditional probabilities for this BN is provided in Supplementary Material, File 6 (software-generated documentation) and File 9 (expressions for generating CPTs).

#### 3.4.1. Nodes representing scenarios and risk metric thresholds

The set of root parent nodes of the BN (Figure 4) is comprised by the main scenario node (“Spat_temp”) and the three risk RQ threshold nodes (Table 4e), along with the substance-specific single-value (function) nodes “HC5” (Table 2). The model can be run by instantiating one or more of the multi-state parent nodes (setting 100% probability for a given state, see Figure 2b). The parent scenario node “Spat_temp” has uniform distribution across the 60 states as the default setting. The uniform distribution is inherited by the two child nodes “RBD” and “Month”. This is also the case for the temporally aggregated scenario node “Season”, while the spatially aggregated scenario node “Region” has a default distribution that reflects the frequency of RBDs (40% Flanders, 60% Wallonia).

A uniform distribution is also used as the initial setting for the three RQ threshold nodes (“Threshold_RQ”, “Threshold_SumRQ”, “Threshold_SumSumRQ”). However, the threshold value 1 is used the default value for model running, reflecting conventional use of RQ for screening-level risk assessment in ecotoxicology.

#### 3.4.2. Nodes in substance-level submodels

For each substance, the set of nodes starting with “Conc” reflects the components of the statistical distributions estimated for the simulated concentrations (PECs), as described in Section 3.1.3. For each substance, the statistical distribution of the log-transformed non-zero PEC values was estimated as a mixture of two normal distributions: λ_1_ × N(µ_1_, σ_1_) + λ_2_ × N(µ_2_, σ_2_), where λ_1_ + λ_2_ = 1. The numbering of the two components was defined by order of the estimated means (µ_1_ < µ_2_). In the BN, the two normal distributions are represented by the nodes “Conc_log_lo” are “Conc_log_hi”, respectively, while the weight λ_2_ is represented by the Boolean node “Conc_log_hi_lambda”. For node “Conc_log_lo”, the CPT-generating expression is the distribution estimated for each spatio-temporal, organised as a chain of 60 “if, then, else” statements (Eq. 1, see example in Table 4a):

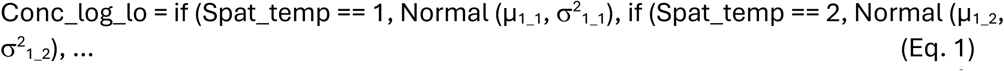

For “Conc_log_hi”, similarly, the expression makes use of the 60 estimated Normal (µ_2_n_, s^2^_2_n_). (Note that the Normal distribution expression in HUGIN uses the variance σ^2^ rather than the standard deviation σ).

The Boolean node governing the weight of the two components (“Conc_log_hi_lambda”) was set to represent the probability of the higher-concentration component (Eq. 2), using the expression Distribution (FALSE, TRUE):

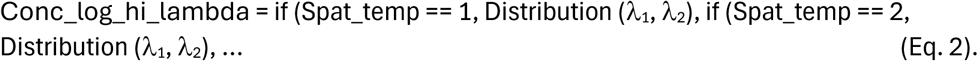

The two components of the log-transformed non-zero concentration are weighted and combined into the node “Conc_log_nz” with a simple “if, then, else” statement (Eq. 3):

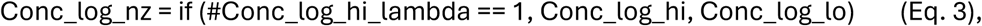

where the expression #[Node] == 1 means TRUE (the second state of the Boolean node). Subsequently this node was back-transformation to natural scale (Eq. 4):

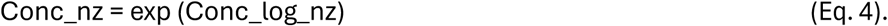

The calculated proportion (p) of zero values for each spatio-temporal unit was represented by the node “Conc_zero_prop” (Eq. 5):

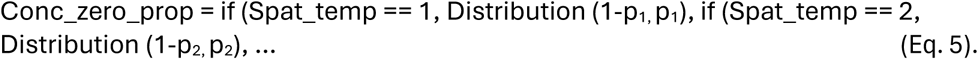

Finally, the expression for the exposure characterisation “Conc” (Eq. 6) combines the probability of zero values with the distribution non-zero values:

Conc = if (#Conc_zero_prop == 1, 0, Conc_nz)

The remaining substance-level nodes (“RQ_unadj”, “RQ” and “RQ_exceeds”) all have simple expression as listed in Table 4a and explained in section 3.2.1.

#### 3.4.3. Nodes in group- and mixture-level submodels

The Concentration Addition method is in implemented in the BN by addition of RQ nodes, in principle corresponding to Eq 6:

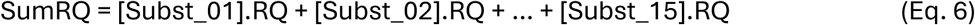

However, to limit the size of the resulting CPT (as explained in Section 3.2), the addition is performed sequentially with intermediate nodes (Table 4b).

The Independent Action method is interpreted as the joint probability of any (i.e. at least one) RQ_n_ exceeds the given threshold (thr.) (Table 1). The following set of equations demonstrates the reasoning. The IA method is implemented with an “or” expression in HUGIN, corresponding to Eq. 7a:

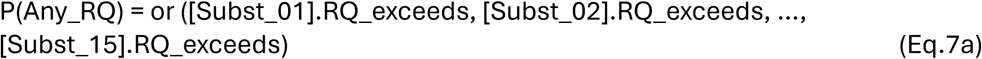

This is equivalent to the following expression with union symbol (∪ = “or”) (Eq. 7b):

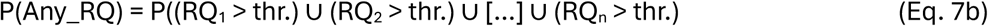

The opposite situation (complementary probability 1-P) is the joint probability that none of the RQs exceed the threshold, in other words that all RQs are below the threshold, which can be expressed with intersection symbol (∩ = “and”) (Eq. 8a):

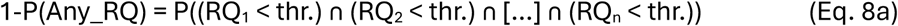

This situation can be expressed by multiplying the probabilities of non-exceedance (Eq. 8b):

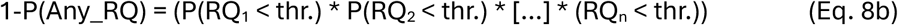

Finally, the expression can be rephrased from the probability of non-exceedance (P(RQ_1_ < thr.)) to the complementary probability of exceeding (1-P(RQ_1_ > thr.)) (Eq. 8c):

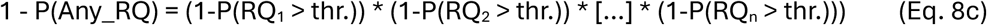

which in turn is equivalent to Eq. 8d:

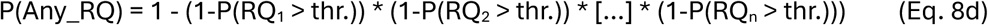

The last expression (Eq. 8d) corresponds to the equation used more specifically in ecotoxicology for combined response to multiple stressors with the Independent Action concept (Eq. 9):

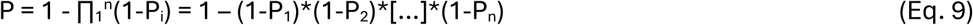

Thus, this final node provides a mixture risk characterisation corresponding to the IA concept: the joint probability of impact from any substance exceeding the given threshold.

The method labelled “CA+IA” (Table 1) combines these two calculation methods by first applying the CA-based calculation across substances within a group, then applying the IA-based calculation across groups within a mixture. In this study, all three methods are applied to the mixture level in order to compare the outcomes, both considering the protectiveness (which method predicts the highest risk) and their ranking of risk-driving substances.

### 3.5. Scenarios and sensitivity analysis

The spatial and temporal extent of the BN developed for this pilot study represents Belgium in year 2012, as described in Section 3.1.1. The spatial resolution is River Basin District, while the temporal resolution is month, while the as described in Section 3.1.3. The set of alternative spatial and temporal scenarios is therefore 5 RBDs x 12 months = 60 spatio-temporal scenarios. The more aggregated versions of spatial and temporal nodes - Region and Season, respectively - will be determined by these settings.

In addition to the spatial and temporal scenarios, a third type of scenarios defined for this BN is the RQ threshold, which have the six alternative states ranging from 0.001 to 100. The BN has been run for all possible 60 x 6 = 360 scenarios, with outputs provided in Supplementary File 10. The assessment reported in this paper will focus on two of the RQ thresholds. (i) Threshold_RQ = 1, for assessing the probability of Conc. ≥ HC5. This corresponds to the criterion PEC > PNEC (Predicted No-Effect Concentration) in lower-tier risk assessments, to screen for substances or cases that need more specific assessments (Duarte et al., 2022). (ii) Threshold_RQ = 0.1, which can be considered an early-warning situation, in particular in the context of mixture risk. This threshold is more arbitrary, but consistent with low-risk thresholds used in comparable works on mixture risk assessment (Dulio et al., 2024).

In the predefined scenarios reported in Supplementary File 10, the same value has been applied to RQ thresholds at all three levels (substance, group, and mixture). However, the model can also be run with different threshold values for the three levels (Threshold_RQ, Threshold_SumRQ and Threshold_SumSumRQ; see Section 3.6).

The sensitivity of mixture risk nodes (see Table 1) to each of the substances and groups was calculated by Value-of-information (VoI) analysis. This method can quantify the potential benefit of additional information in the face of uncertainty, and can aid the decision on allocation of resources between obtaining new information and improving management actions (Mäntyniemi et al., 2009). The method quantifies the VoI as reduction of entropy (a measure of uncertainty) for a given node (e.g. a substance) in isolation, averaging the values of other nodes. The outcome of the sensitivity analysis is presented in Section 4.2 (Ranking of risk drivers).

### 3.6. Online user interface

In addition to the full model specification for the BN (Supplementary File 5), an online user interface is provided in a publicly available web site: https://encore.hugin.com/models/Pilot. The purpose of this interface is to enable quick access to the main parts of the model, and to allow users to run the model with default settings and inspect the outcome via the main input and output nodes. The user interface with default settings is displayed in Supplementary File 7.

The seven available input nodes correspond to the scenarios (Section 3.5): Spatial settings (RBD or Region), Temporal settings (Month or Season), and RQ thresholds (for Substance, Group and Mixture).

Considering the temporal scenarios, the model is meant to be run from either Month or from Season. For example, if Season is set to Spring, then the Month node will get 33.33% probability for each of the states March, April and May. In case evidence is given for both levels, inconsistent evidence will result in an error message. Likewise, the spatial scenarios are meant to be run from either RBD or from Region.

The RQ threshold nodes can be set to any of the alternative states (from 0.001 to 100). The thresholds can be set independent for the three levels (Substance, Group and Mixture). However, the assessments in this paper only considers the cases with RQ thresholds 0.1 and 1, and only scenarios with the same threshold applied to all three levels.

The output nodes available through the user interface are the 21 Boolean nodes representing the probability of threshold exceedance (RQ > Threshold_RQ), for the 15 substances, the 3 substance groups and the 3 mixture risk calculation methods.

## 4. Results

The results are presented in two subsections, according the main purposes of this study: (1) to assess the model’s alternative mixture risk calculation methods across a set of scenarios, and (2) to assess the BN’s performance in ranking of risk-driving substances. The BN model output referred to below is available in Supplementary File 10.

### 4.1. Risk characterisations

In this study, “risk” is defined by two components (Table 1): both the risk metric (RQ) and the probability that this metric exceeds a given threshold (Threshold_RQ). The results reported here focus on two selected RQ thresholds, representing screening assessment (Threshold_RQ = 1) and early warning assessment (Threshold_RQ = 0.1).

#### 4.1.1. Substance-level risk

For each substance, the risk metric (RQ) is quantified as a probability distribution across the 12 intervals, for each RBD and month. As an example, the RQ distributions for the RBD MAAS is illustrated in Figure 5, while the corresponding figures for the remaining four RBDs are given in Supplementary File 11. To aid the interpretation, the length of the bar segments with red and orange colour combined represent the probability of RQ > 1, while the length of the bar segments with red, orange and yellow colours combined represent the probability of RQ > 0.1. Bar segments with green and blue colours represent lower RQ thresholds (0.01 and 0.001, respectively). While these risk metric levels are negligible when considering risk of individual substances, they still have the potential to contribute to group-level or mixture risk.

**Figure 5.**
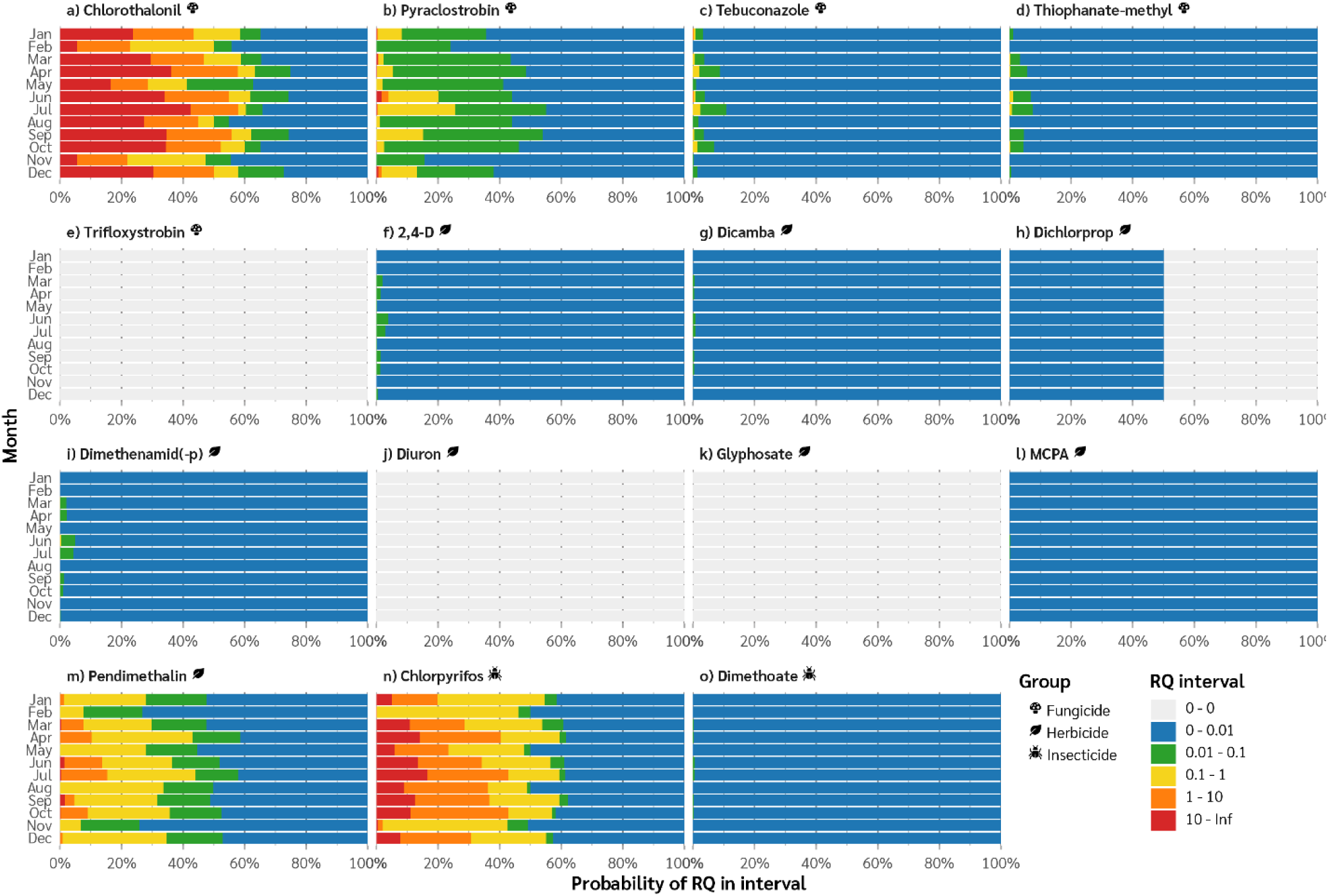
Substance-level risk: Predicted probability distribution of RQ across six intervals (see legend), specific for each river basin district (RBD), month and substance (a-o). The respective substance groups (Fungicide, Herbicide, Insecticide) are indicated by icons. Results are shown for the river basin district MAAS (see Figure 1).

Visual inspection of Figure 5 suggests that for RBD MAAS, only three substances have RQ > 1 for substantial parts of the year: the fungicide Chlorothanlonil (Figure 5a), the insecticide Chlorpyrifos (Figure 5n) and the herbicide Pendimethalin (Figure 5m). In addition, the fungicide Pyraclostrobin (Figure 5b) shows low probability of exceedance of 1 in a few months. Thus, these 3-4 substances are the main drivers of the risk patterns observed for the respective substance groups. These are all with substances relatively high toxicity (HC5 values in ranging from 0.0008966 to 0.3409 µg/L; Table 2).

For three of the substances (Trifloxystrobin, Diuron and Glyphosate), the simulated concentration was either zero or below the lower limit of 10^-9^ µg/L, resulting in zero risk for the RBD MAAS. The remaining 8 substances mostly had RQ levels below 0.01. While these risk levels can be negligible in the given setting with 15 substances, such contributions can still matter in real cases with a considerably higher number of substances.

Most of the substances with non-zero risk estimates displayed a conspicuous seasonal pattern, with peak risk for every two to three months. For example, for Chlorothalonil (Figure 5a), peak probabilities of RQ > 0.1 can be seen in the months April, July, September, and December. These temporal patterns in risk reflect the patterns in PECs and risk reported by Mentzel et al. (2026) for Pilot 2. The explanation is related to the assumed pesticide application scheme used in the process-based simulation, in combination with seasonal climatic and hydrological processes. An evaluation of whether these particular patterns are realistic is beyond the scope of this paper. However, the temporal patterns of risk metrics represent variation in RQ distributions that should be addressed in the sensitivity analysis.

#### 4.1.2. Group-level risk

The risk characterisation at the group level is displayed for the CA method, i.e. the probability distribution of SumRQ (Figure 6), for each substance group (vertical panels) and RBD (horizontal panels) as well as for each month (horizontal bars, as in Figure 5). In addition, the group-level risk characterisations were calculated with the IA method, i.e. the joint probability of RQ > Threshold_RQ for all substances within a group (not displayed but available in Supplementary File 10). The spatio-temporal patterns in the CA risk characterisation largely reflected those of the dominant risk-driving substances in each group. For example, considering fungicides in the RBD MAAS (Figure 6j), peak probabilities of RQ > 1 (red + orange bar segments) can be observed for the months April, July, September and December, corresponding to the peak risk months for the fungicide Chlorothalonil (Figure 5a).

**Figure 6.**
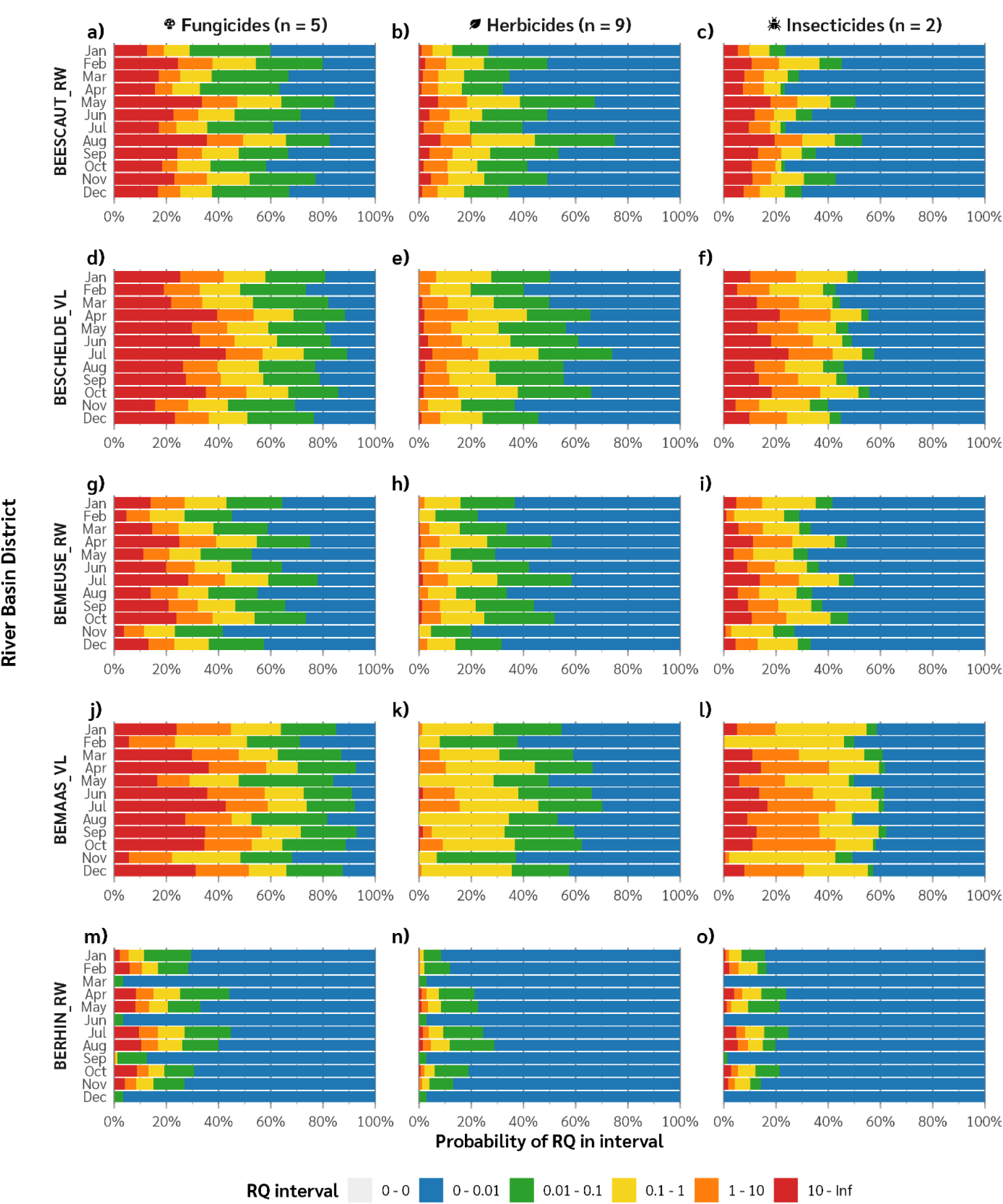
Group-level risk: Predicted probability distribution of RQ across six intervals (see legend), specific for each river basin district (horizontal panels), month and substance group (vertical panels).

Visual inspection of the overall risk patterns at the group level showed that the highest risks (e.g., the longest red and orange bar segments) were posed by fungicides, followed by insecticides. This general pattern was consistent for all RBDs, although the temporal patterns could vary across RBDs. For example, insecticide risk peaked in April and July for RBDs MAAS, MEUSE and RHIN, but in May and August for ESCAUT. The explanation is a combination of application (timing, crop types in the SC’s, and amount of agricultural land in the SC) and hydrology (emission to water, water flow between SC’s, partitioning and degradation) (Mentzel et al., 2026).

Comparing the risk patterns across RBDs, the first four RBDs (Figure 6a-l) had comparable risk levels, while RHIN stood out with lower risk (Figure 6m-o). The more peripheral geographic location of RHIN in the southern part of Belgium (Figure 6) could explain why the subcatchments in this RBD had lower PECs and consequently RQ values. Another observation from Figure 6a-l was that RBDs in the region Vlaanderen (SCHELDE and MAAS) displayed slightly higher risk than their counterparts in Wallonia (ESCAUT and MEUSE).

Considering the assignment of substances to groups for risk assessment, various approaches have been proposed in the literature, including cumulative assessment groups (EFSA Scientific Committee et al., 2021), consensus mode of action groups (Kienzler et al., 2019) and common kinetic groups (Braeuning et al., 2022). In the current pilot study, the group assignment reflects a pragmatic aim: obtaining a balanced allocation of groups per mixture and substances per groups, to enable a reasonable comparison of the CA+IA calculation methods with the two other alternatives. For future development and expansion of this BN, the group assignments can easily be adapted to different definitions of substance groups, given the modular and flexible structure of the BN (Figure 4).

#### 4.1.3. Mixture-level risk

The risk characterisation for the mixture level is displayed for all three calculation methods (Figure 7), using the RBD MAAS as an example. Risk characterisation with the CA method is displayed as the discrete probability distribution of SumSumRQ (Figure 7a). The layout is similar as for substance level (Figure 5) and group level (Figure 6), but with shaded colours, to highlight and aid the probabilistic risk interpretation. For the default risk screening setting Threshold_RQ = 1 (upper panel), the probabilistic interpretation of mixture risk with the CA method is the probability of SumSumRQ > 1, corresponding to the total length of the red and orange bars (Figure 7a). Likewise, for the early-warning setting Threshold_RQ = 0.1 (lower panel), mixture risk with the CA methods corresponds to the total length of the red, orange and yellow bars (Figure 7d).

**Figure 7.**
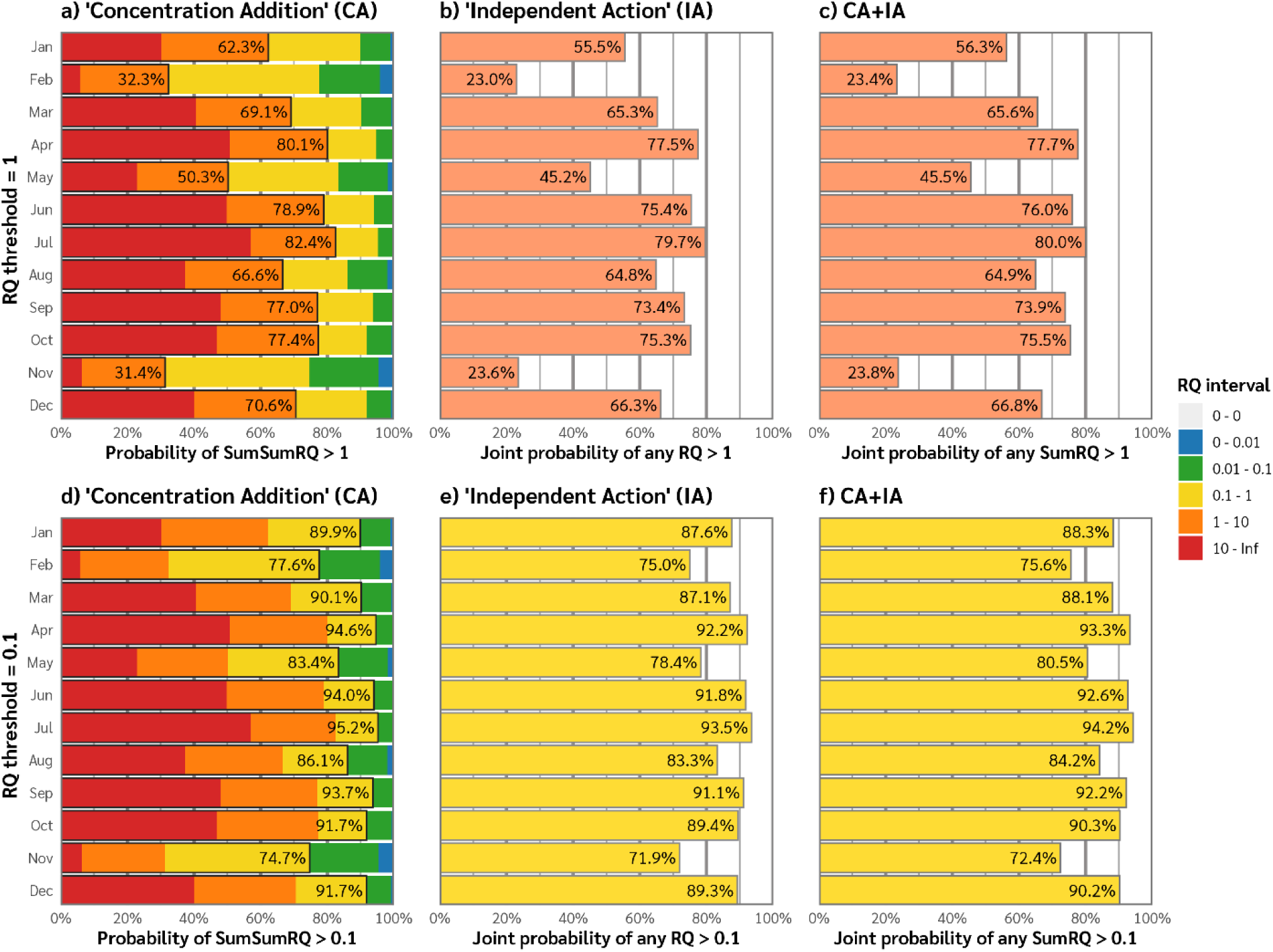
Mixture-level risk characterisation: Predicted probability of the given risk metric exceeding the given RQ threshold (RBD = BEMAAS_VL). Upper panel (a-c): RQ Threshold = 1; Lower panel (d-f): RQ Threshold = 0.1. For explanation to the three methods for mixture risk calculation, see Table 1). For the plots based on CA calculation only (a and d), the stacked bars show the intervals of SumSumRQ, while the outlined bar segments represent the probability of SumSumRQ exceeding the given threshold. For the plots involving joint probability calculation (IA and CA+IA), the stacked bar display is not possible.

Mixture risk characterisation with the IA method (Figure 7c) shows the joint probability of RQ > 1 for any of the 15 substances, using the expression represented in Eq. 9 (see Table 1). In comparison to the CA method (Figure 7a), the IA method gave very similar risk patterns across the months. However, the IA calculation consistently resulted in slightly lower risk. For example, considering mixture risk predictions for the RBD MAAS (Figure 7), IA-based predictions ranged 23.0% - 79.4% across the 12 months, while the CA-based predictions ranged 31.4% - 82.4%. (Values for other RBDs can be found in Supplementary File 10).

The combined method, as displayed in Figure 7b, shows the joint probability that SumRQ > 1 for at least one of the substance groups in MAAS (Figure 6j-l). Visualisation with stacked bars (as in Figure 7a) is not possible when the SumRQ values have been combined with the joint probability calculation. However, Eqs. 7-8 can help interpret the combined-method risk in connection to the substance-level bars. For example, for MAAS in May, there was 45.5% joint probability of one or more SumRQ exceeding the value 1.

The combined method CA+IA consistently provided risk values in between those from CA and IA, for all months and RBDs. This outcome could be expected, given that the methods integrate calculations from the two other methods. Nevertheless, this outcome suggests that the CA+IA method is a reliable method which can be expected to give less extreme risk predictions than either CA or IA, and to more robust for example to outliers.

In the “early warning situation” with RQ threshold = 0.1 (Figure 7 lower panel), the predicted mixture risk for MAAS was considerably higher (71.9% - 94.6%) across all months, compared to the default situation (RQ threshold = 1). In general, lowering the RQ threshold from the screening level (1) to the early warning level (0.1) resulted in considerable increases in predicted mixture risk, typically from 40-50% (Figure 7 upper panel) to 80-90% (Figure 7 lower panel). In some cases, the increase in predicted mixture risk was even more extreme, such as from <35% to >75% (e.g. February and October). Which RQ threshold is the most appropriate for risk assessment and management in different situations is another discussion, and beyond the scope of this paper. The higher risk patterns in the “early warning situation” appeared less variable across the months and across the calculation methods (vertical panels). For the temporal patterns in mixture risk, the lower RQ threshold mostly resulted in the same ranking of months as the default RQ threshold. Considering the alternative methods, inspection of the numbers (Supplementary File 10) revealed the same type of patterns as for the “screening situation” (Figure 7 upper panel): the IA method predicted consistently slightly lower mixture risk than the CA method, with the combined CA+IA method producing a compromise.

### 4.2. Ranking of risk drivers

The substance groups have been ranked as risk drivers according to three different types of metrics, based on the relevant RQ-related nodes, under specific scenarios (Table 5a): (1) the posterior mean value (i.e. the expected value) of SumRQ; (2) the posterior probability of both group-level risk calculations (SumRQ_exceeds and Any_RQ_exceeds) given both thresholds (1 and 0.1); and (3) the sensitivity of the mixture risk (CA+IA method) to the SumRQ of each group. Likewise, the individual substances have been ranked by the same three type of metrics at the substance level; results for the top 5 risk-driving substances are displayed in (Table 5b). In both cases (Table 5a and b), the default rank is based on the mean (Sum)RQ. All mentioned mean values and probabilities are available in Supplementary File 10, while the sensitivity scores are available in Supplementary File 12.

**Table 5.** Ranking of groups (a) and substances (b) according to their contribution to mixture risk, and by sensitivity analysis. “Mixture risk” is here defined by the variable “Any SumRQ exceeds?” (method CA+IA; Table 1), with RQ threshold (“Thrs.”) set to 1 or 0.1 (a) Group level. (No selection of months). For each RBD, the three substance groups are ranked decreasing value of by five risk metrics: “Mean SumRQ” (corresponding to traditional CA), “SumRQ exceeds 1” (probabilistic CA, screening situation), “Any RQ exceeds 1” (IA, screening situation), “SumRQ exceeds 0.1” (probabilistic CA, early warning situation) and “Any RQ exceeds 0.1” (IA, early warning situation). The sensitivity scores are obtained by value-of-information analysis, calculated as the mutual information of the hypothesis variable (mixture risk) to each of the information variables (SumRQ for each group). (b) Substance level, showing results for the five substances with highest risk, for three selected months (Feb, May, Aug). Otherwise, the calculations correspond to those for group level (a). Asterisks (*) highlight any cases where the ranking deviates from the default ranking based on MeanSumRQ.

| River Basin District | Month | Substance Group | Mean SumRQ |  | SumRQ exceeds 1? |  | Any RQ exceeds 1? |  | SumRQ exceeds 0.1? |  | Any RQ exceeds 0.1? |  | Mixture risk (CA+IA) sensitivity to SumRQ |  |
| --- | --- | --- | --- | --- | --- | --- | --- | --- | --- | --- | --- | --- | --- | --- |
|  |  |  | Rank | Value | Rank | Value | Rank | Value | Rank | Value | Rank | Value | Thrs. = 1 | Thrs. = 0.1 |
| ESCAUT | All | Fungi | 1 | 6.48 | 1 | 31.3 % | 1 | 30.8 % | 1 | 44.9 % | 1 | 43.2 % | 0.29 | 0.25 |
| ESCAUT | All | Insec | 2 | 3.64 | 2 | 19.2 % | 2 | 19.2 % | 2 | 28.3 % | 2 | 28.2 % | 0.16 | 0.13 |
| ESCAUT | All | Herbi | 3 | 1.28 | 3 | 11.0 % | 3 | 10.9 % | 3 | 24.2 % | 3 | 23.3 % | 0.08 | 0.11 |
| SCHELDE | All | Fungi | 1 | 8.32 | 1 | 42.0 % | 1 | 41.2 % | 1 | 58.0 % | 1 | 55.9 % | 0.29 | 0.18 |
| SCHELDE | All | Insec | 2 | 4.60 | 2 | 28.8 % | 2 | 28.8 % | 2 | 44.0 % | 2 | 43.9 % | 0.17 | 0.11 |
| SCHELDE | All | Herbi | 3 | 0.94 | 3 | 11.7 % | 3 | 11.7 % | 3 | 30.3 % | 3 | 29.3 % | 0.06 | 0.07 |
| MEUSE | All | Fungi | 1 | 4.71 | 1 | 27.3 % | 1 | 26.8 % | 1 | 41.3 % | 1 | 40.3 % | 0.33 | 0.25 |
| MEUSE | All | Insec | 2 | 2.35 | 2 | 16.1 % | 2 | 16.1 % | 2 | 31.8 % | 2 | 31.8 % | 0.17 | 0.17 |
| MEUSE | All | Herbi | 3 | 0.36 | 3 | 4.8 % | 3 | 4.8 % | 3 | 17.2 % | 3 | 16.7 % | 0.04 | 0.08 |
| MAAS | All | Fungi | 1 | 7.68 | 1 | 45.6 % | 1 | 45.1 % | 1 | 62.1 % | 1 | 59.5 % | 0.35 | 0.14 |
| MAAS | All | Insec | 2 | 3.32 | 2 | 28.2 % | 2 | 28.2 % | 2 | 53.4 % | 2 | 53.4 % | 0.17 | 0.11 |
| MAAS | All | Herbi | 3 | 0.36 | 3 | 5.3 % | 3 | 5.3 % | 3 | 30.9 % | 3 | 30.0 % | 0.03 | 0.05 |
| RHIN | All | Fungi | 1 | 1.42 | 1 | 8.4 % | 1 | 8.3 % | 1 | 13.5 % | 1 | 13.3 % | 0.21 | 0.24 |
| RHIN | All | Insec | 2 | 0.64 | 2 | 3.7 % | 2 | 3.7 % | 2 | 8.0 % | 2 | 8.0 % | 0.08 | 0.13 |
| RHIN | All | Herbi | 3 | 0.19 | 3 | 1.5 % | 3 | 1.5 % | 3 | 4.3 % | 3 | 4.2 % | 0.03 | 0.07 |

(b)
| River Basin District | Month | Substance (Group) | Mean RQ |  | RQ exceeds 1? |  | RQ exceeds 0.1? |  | Mixture risk (CA+IA) sensitivity to RQ |  |
| --- | --- | --- | --- | --- | --- | --- | --- | --- | --- | --- |
|  |  |  | Rank | Value | Rank | Value | Rank | Value | Thrs. = 1 | Thrs. = 0.1 |
| MAAS | Feb | Chloro (Fungi) | 1 | 1.86 | 1 | 22.9 % | 1 | 50 % | 0.52 | 0.21 |
| MAAS | Feb | Chlorp (Insec) | 2 | 0.25 | 2 | 0.2 % | 2 | 46 % | 0.00 | 0.18 |
| MAAS | Feb | Pendim (Herbi) | 3 | 0.02 | 3 | 0.0 % | 3 | 8 % | 0.00 | 0.02 |
| MAAS | Feb | Pyracl (Fungi) | 4 | 0.01 | 4 | 0.0 % | 4 | 0 % | 0.00 | 0.00 |
| MAAS | Feb | Tebuco (Fungi) | 5 | 0.00 | 5 | 0.0 % | 5 | 0 % | 0.00 | 0.00 |
| MAAS | May | Chloro (Fungi)* | 1 | 4.94 | 1 | 28 % | *2 | 41 % | 0.30 | 0.12 |
| MAAS | May | Chlorp (Insec)* | 2 | 2.32 | 2 | 23 % | *1 | 48 % | 0.23 | 0.15 |
| MAAS | May | Pendim (Herbi) | 3 | 0.07 | 3 | 0 % | 3 | 28 % | 0.00 | 0.07 |
| MAAS | May | Pyracl (Fungi) | 4 | 0.03 | 4 | 0 % | 4 | 2 % | 0.00 | 0.01 |
| MAAS | May | Tebuco (Fungi) | 5 | 0.00 | 5 | 0 % | 5 | 0 % | 0.00 | 0.00 |
| MAAS | Aug | Chloro (Fungi) | 1 | 7.48 | 1 | 45 % | 1 | 50 % | 0.29 | 0.13 |
| MAAS | Aug | Chlorp (Insec) | 2 | 3.47 | 2 | 36 % | 2 | 49 % | 0.21 | 0.12 |
| MAAS | Aug | Pendim (Herbi) | 3 | 0.08 | 3 | 0 % | 3 | 34 % | 0.00 | 0.07 |
| MAAS | Aug | Pyracl (Fungi) | 4 | 0.03 | 4 | 0 % | 4 | 1 % | 0.00 | 0.00 |
| MAAS | Aug | Tebuco (Fungi) | 5 | 0.00 | 5 | 0 % | 5 | 0 % | 0.00 | 0.00 |

#### 4.2.1. Ranking of groups

The list of substance groups ranked by decreasing mean SumRQ (Table 5a) is based on values for all months (i.e., no evidence is set for Month). This overview shows consistent ranking of groups by all three main types of metrics: fungicides posing the highest risk, followed by insecticides. The probabilities in the CA-based column “SumRQ exceeds 1?” are consistently slightly higher (or equal to) the probabilities in the IA-based counterpart (“Any RQ exceeds 1?”). The same is true for the next two columns with RQ threshold 0.1.

The sensitivity of the mixture risk node “Any_SumRQ_exceeds” (representing the CA+IA method) is usually higher for RQ threshold 1 than for 0.1, for all RBDs except RHIN. The higher sensitivity given RQ threshold 1 can reflect the higher variability of risk across months with this condition, as shown for the RBD MAAS (compare the high variability of bars in Figure 7b with the lower variability in Figure 7e). For RQ threshold 1, the probability of exceedance (i.e. the right end of the orange bar, Figure 6j-k) varies across the groups, spanning from typically 30-50% for fungicides, via 20-40% for insecticides and 0-10% for herbicides. Here, the change in SumRQ for any of the groups can have a considerable impact on the overall risk (i.e. the probability of any SumRQ exceeding 1). For RQ threshold 0.1, in contrast, all three groups have shown relatively high probability of exceedance (right end of the yellow bar, Figure 6j-k): typically 60-70% for fungicides, 50-60% for insecticides and 30-40% for herbicides. Hence, reduction in SumRQ for any of the groups may have lower influence on the mixture risk in the “early warning” situation.

#### 4.2.2. Ranking of substances

To investigate in more detail the BN’s ranking of the top 5 risk-driving substances, Table 5a is zoomed in to an example RBD (MAAS) and to three selected months (February, May and August). The default ranking by Mean RQ is the same for all three months (as well as for the remaining months; Supplementary File 12): The mixture risk (CA+IA method) is clearly dominated by Chlorothalonil (fungicide), followed by Chlorpyrifos (insecticide). The next three substances have considerably lower influence: Pendimethalin (herbicide), Pyraclostrobin and Tebuconazole (both fungicides). The ranking in the column “RQ exceeds 1?” follows the same pattern as the Mean RQ, while the ranking by “RQ exceeds 0.1?” shows a deviation: for May, the insecticide Chlorpyrifos is ranked higher (48% probability) than the fungicide Chlorothalonil (41% probability), although Chlorothalonil has the higher mean (RQ 4.94 vs. 2.32). The swapped ranking of Chlorothalonil and Chlorpyrifos is reflected in the sensitivity of the mixture risk under these conditions (0.12 and 0.15, respectively).

## 5. Discussion

### 5.1. Mixture risk modelling for environmental assessment and management

The purpose of this study was to develop a Bayesian network model for providing probabilistic risk characterisation for chemical mixtures, based RQ probability distributions obtained from simulated exposure data. In this way, the spatial and temporal variability of chemical exposure of the pilot study could be incorporated in the probabilistic risk characterisation. Comparing the alternative methods for probabilistic mixture risk considered in this study, the overall differences in predicted mixture risk were only marginal (Figure 7). Nevertheless, the IA-based mixture risk characterisation (Any_RQ_exceeds) generally provided slightly lower probability of exceedance than the CA-based method. The IA-based risk characterisation can therefore be considered slightly less stringent or protective than the CA-based risk characterisation. The combined CA+IA method is the alternative that best represents our mechanistic understanding of cumulative risk. This method, which consistently provided a compromise between the two traditional methods, can also be expected to be more robust towards extreme values and uncertainties.

The CA and IA concepts have traditionally been used in ecotoxicology in a more narrow sense, with response variables corresponding to measured values from toxicity testing. For example, CA and IA models can be used to predict a demographic response (e.g. mortality) of test organisms to given mixture levels. Conversely, such models can be used to derive the lowest concentration levels that cause a given response level (e.g. 50% lethality (LC50), or other type of effect (EC50)). Here we have used the CA and IA concepts in a more generic sense, to predict either the probability of summed RQs exceeding a threshold; or the joint probability of one or more RQs exceeding the threshold. Nevertheless, the general pattern of risk predictions found in our study is consistent with typical with findings in the literature: the CA methods tends to predict slightly higher mixture toxicity than the IA. For example, Backhaus et al. (2004) studied

the combined effects of six toxicants with dissimilar mode of action on the carbon fixation of natural algal communities. As expected given this type of mixture (with dissimilar substances), they found that an IA model predicted the experimental EC50 accurately, while CA overestimated the toxicity slightly (i.e., underestimated the EC50) (Backhaus et al., 2004).

Conversely, in an experimental study with eight similarly acting herbicides, (Junghans et al., 2003) found that a CA model accurately predicted the combined effect on algal reproduction while IA underestimated the mixture toxicity. Similar trends were revealed in studies with larger mixtures (25 pesticides), representing realistic scenarios in agricultural run-off waters revealed similar trends (Junghans et al., 2006), with IA giving slightly higher lower toxicity estimates (higher EC50 values) than CA. The authors concluded that CA provides a precautionary but not overprotective approach for combined effect predictions of pesticide mixtures under realistic exposure scenarios.

Since the CA approach is both computationally simpler and more stringent compared to IA, CA has been favoured as a pragmatic and precautious choice for mixture risk assessment in regulatory review and guidance documents (e.g., Bopp et al., 2015; EFSA Scientific Committee et al., 2021; SCHER et al., 2012). However, these recommendations are largely based on a comparison between the CA and IA methods, while omitting the third option - a combination. Early works on mixture toxicity with combined experimental and modelling studies have demonstrated that a combination of CA and IA models can outperform both CA and IA (Sigurnjak Bureš et al., 2021).

One example is the two-stage prediction (TSP) model tested for effects realistic aquatic pesticide mixtures on algal growth (Junghans, 2004). The TSP technique applied CA calculation for subgroups of substances with anticipated similar modes of action, followed by IA calculation (referred to as response addition) for independently acting groups. Similarly, in a study of mixture toxicity of different nitrobenzenes on algal reproduction, calculation according to the TSP performed better than CA (Altenburger et al., 2005). In this case, the IA-type predictions were derived using the calculated effects of the substances applied singly at certain concentrations, using their Hill or Weibull function. The superior performance of the TSP model has also been supported by later validation studies (Mo et al., 2017).

Combined CA+IA type approaches have also been used more generally, with response variables representing higher levels of biological organisation such as species assemblages. Based on the concept of msPAF (multi-stressor Potentially Affected Fraction of species), de Zwart and Posthuma (2005) advocated the use of a two-step, mixed-model approach: mixture toxicity for individual modes of action to be evaluated with the CA model, and the toxicities of different modes of action are combined using the Response Addition model (equivalent to IA). A corresponding two-step approach for mixture risk calculation has been implemented in PERPEST, a tool for probabilistic prediction of pesticide risk to major species groups (Van den Brink et al., 2006).

More recently, a review on mixture risk assessment concluded that the conventional models (CA and IA) are still most commonly used, even though the combined models (CA+IA types) are more accurate compared to conventional ones (Sigurnjak Bureš et al., 2021). One explanation can be that the application of a combined CA+IA model requires more data, at least compared to CA alone. This is true for the use of IA in the narrow sense, which require a concentration-response relationship from toxicity testing. However, the principles of joint probability for independent events can also be applied more generally in risk assessment, for example, for combining PAFs (potentially affected fractions of species) for multiple pesticides (Neelamraju et al., 2025). When IA is applied more generally to response variables expressed as proportions, and more types of information can be involved, the data requirements are not necessarily higher for IA than for CA, as demonstrated here. Since the early TSP model (Junghans, 2004), models based on the CA+IA principle have been further developed in various directions: for example, including fuzzy membership functions (Wang et al., 2009), multiple linear regression (Qin et al., 2011), and QSAR (quantitative structure-activity relationships). The model presented here, however, is the first published implementation of a CA+IA type mixture risk model in a BN, to our knowledge.

Looking beyond ecotoxicology, the issue of multiple stressors and combined effects on biological systems is a main topic also in ecology, for example in quality assessment for aquatic ecosystems. As pointed out by Schäfer et al. (2023), these two related research areas face similar challenges to predict joint effects of multiple stressors, but still tend to use different tools and methods. While the concept of independent action is not commonly used in ecology, an analogous calculation method was used by Schinegger et al. (2016) for assessing the combined effects of multiple physico-chemical stressors on fish communities by Ecological Quality Ratios (EQR, scale 0-1). A stressor-specific EQR can be understood as the probability of an ecosystem affected by this stressor to achieve the highest possible assessment value (EQR = 1). Assuming that different stressors have independent actions, the combined effects could therefore be predicted as the joint probability of the involved single stressors, i.e. the product of their EQRs (Schinegger et al., 2016).

Another challenge in mixture risk assessment is interactions, i.e., synergistic or antagonistic effects. Interactions are often addressed both in the context of chemical mixtures (e.g., Hernandez et al., 2019) and multiple stressors (e.g., Schinegger et al., 2016). However, reviews of the available data indicate that synergisms and antagonisms very rarely lead to a deviation from the CA-predicted mixture toxicity (Backhaus, 2016). Furthermore, it has been argued that both multiple stressor and chemical mixture research are too focused on interactions (Schäfer et al., 2023). In line with these statements, we have not accounted for potential interactions among the effects of pesticides in the BN model presented here.

### 5.2. Evaluation of the BN approach to mixture risk modelling

Bayesian network methodology has been used extensively for environmental impact and risk assessments (Kaikkonen et al., 2020; Landis, 2021), in particular for assessments of aquatic ecosystems and water quality (Borsuk et al., 2004). BNs for water quality assessment have often focused physico-chemical factors related to eutrophication and other pressures, sometimes in combination with climate change (Adams et al., 2023; Molina-Navarro et al., 2020). More recently, BN applications have focussed on specific chemical stressors or groups such as pesticides (Piffady et al., 2021), pharmaceuticals (Welch et al., 2024) and mercury (Jermilova et al., 2025).

Modelling approaches to risk assessment have traditionally divided into two main types: (1) deterministic, e.g., risk characterisation by a single-value risk score; or (2) probabilistic, e.g., risk characterisation involving a probability distribution, or the probability of a given outcome. For BNs, however, this dichotomy does not strictly apply (Moe et al., 2022): the BN described here is essentially a deterministic model, while offering probabilistic risk characterisation. Compared to other process-based models, this dual property of BN models provide additional opportunities for inference, as will be elaborated below.

Modelling of environmental systems with BNs has well-known constraints, such as the need for discretisation of variables. The BN software used in this study allows child nodes to be truly continuous variables (Gaussian; see Mentzel et al., 2026), but not for parent or intermediate nodes. (While BNs constructed in some softwares can appear to have continuous nodes in such locations, these are usually approximations with high resolution of intervals). The challenges associated with discretisation of variables in this study are adressed in Section 3.3, along with the chosen solutions. To assess the robustness of the BN’s model prediction to these discretisations, we could explore alternative discretisations schemes and compare the resulting ranking of substances. However, since the mixture risk in this pilot study turned out to be clearly dominated by a few substances, it is not likely that changes in discretisation would have much effect on the ranking.

The use of joint probability calculation (based on the IA concept) in this BN implies assumptions of independence for both exposure and effects of substances in different groups. Whether such assumptions are reasonable will have to be evaluated from case to case. When the BN is later expanded to incorporate upstream processes such as chemical emission and transport, then correlations (e.g., pesticides applied to at the same time) can potentially be handled in the upstream components of the BN.

In this study, environmental risk was quantified by RQs as a simple risk metric, compared to more ecologically relevant endpoints such as SSDs (de Zwart and Posthuma, 2005) or species groups (Van den Brink et al., 2006). The use of SumRQ or equivalent approaches are common for pesticides as well as for other types of substances, such as pharmaceuticals (Duarte et al., 2022). Nevertheless, the summing of RQs has conceptual flaws and have been reasonably criticised. For example, (Topping et al., 2026) questioned the use of single-point RQs in combination with a mixture allocation factor. In our study, the use of RQs was a pragmatic choice to limit the complexity of the risk characterisation component of the BN, while developing the multi-level structure for propagating information within and across groups. The RQ probability distributions in this study were calculated as a ratio of a distribution (Conc.) to a single value (HC5). The denominator (HC5) could be refined to represent a distribution as well, for example an SSD (while considering the role of precautionary factors), as described for a case study in Norway (Mentzel et al., 2022). The BN structure can also be further developed to incorporate more meaningful ecological responses than RQs. For example, for pesticides, the case-based reasoning system PERPEST offers probabilistic effects characterisation for aquatic species groups, which can be linked from to the probabilistic exposure characterisation of the current BN. Using a case study from rice fields in Spain, Mentzel et al. (2024) demonstrated how a BN can be used to combine process-based model predictions with information from PERPEST, in the context of climate change scenarios. The case study was highlighted as a promising approach for connecting chemical pollution with effects on biodiversity by the European Joint Research Centre (Baccaro et al., 2025). By including species groups as assessment endpoints in the BN, we can also better meet requirements for seperate assessments of different trophic levels or functional groups.

The pilot study reported here has demonstrated several strengths of BN methodology for mixture risk characterisation. (1) The calculation of IA-based mixture risk, i.e. joint probabilities for independent events, can easily be implemented in BNs with OR expressions (or manually by conditional probability tables). (2) The modular nature of OOBNs enables a flexible and transparent construction of the multi-level structure with substances, groups and mixtures. (3) The quick computation time of the BN allows for efficient exploration of alternative scenarios, for example, alternative RQ thresholds to represent higher or level degrees of precautionary settings. (4) Because the BN uses deterministic and fully traceable calculations, sensitivity scores can be derived directly from the parameterised BN (without the need for simulations); these scores can support the ranking and prioritisation of risk-driving substances. (5) The graphical structure of the BN is helpful for visualising and communicating complex models structures, in particular for OOBNs. (6) The BN can be extended and further developed, for example, to include chemical emission components and/or more ecologically relevant assessment endpoints.

For practical use of BN in its current form, parameterised for the given pilot study, the main constraint may lie in the resolution of continuous variables. Moreover, the spatial resolution may be too coarse for supporting river basin management and decisions in practice. Therefore, the most useful application could be for ranking of substances by their influence on the mixture risk, rather than the exact predicted risk value (the probability of exceedance). Moreover, and after expansion to larger spatial scales, the BN can be used for comparison of relative risk across different regions, periods and other setting, as demonstrated by the Bayesian Network Relative Risk Model (Landis, 2021). Such a scenario-based approach can be in line with the recommendation of (Holmes et al., 2017): “to determine whether mixtures of chemicals pose risks over and above any identified using existing approaches for single chemicals, how often and to what magnitude, and ultimately which mixtures (and dominant chemicals) cause greatest concern”.

The current pilot study was limited to one country (represented by 5 river basin districts) and one year (divided into 12 months), as a starting point. Based on the currently available output for 23,000 subcatchments from the ENCORE exposure model (Vlaeminck et al., 2026), this type of BN can in principle be constructed for more than 1000 substances in 42 countries divided in to RBDs (or other spatial units) for 10 years (2004-2013). Furthermore, all simulations are available in 100 stochastic replicates, representing variability in use volumes (REACH chemicals and pharmaceuticals), environmental release factors (REACH chemicals), and application rates (pesticides). The variation across the replicated simulations could also be accounted for in the BN as an emulation model, in addition to the spatial and temporal variability which is currently incorporated. The open access to all ENCORE exposure model simulations (Vlaeminck et al., 2026), combined with the full BN model specification in this paper, will allow other researchers to further explore and develop the approach presented here. Moreover, R scripts for efficient construction of BN submodels for substances and groups can be shared upon request.

If the BN is adapted to different study areas, the spatial data structure should be reconsidered: in some countries the RBD (or RBD subunit) structure might have too low resolution for giving useful predictions. However, the BN methodology also allows spatial refinement by combining probability distributions with new evidence (Bayesian updating; see example in Mentzel et al., 2026). This way, for a given water body and a given chemical substance, new monitoring data from the water body (evidence) can be combined with the estimated concentration for the respective RBD (prior probability distribution) to obtain a refined concentration estimate for the water body (posterior probability distribution).

The comparison of the three methods for mixture risk calculation showed only small differences (Table 5, Figure 7). Therefore, based on this pilot study, one may question the utility of a tool for mixture risk calculation that enables all three methods. However, the high similarity of outcomes in this study can partly be explained by the fact that each substance group was dominated by one substance as the risk driver (Figure 5). An extended BN that includes a higher number of substances and substance groups can provide more insights into the behaviour of the three methods for probabilistic mixture risk calculation, under different scenarios. For example, it can be hypothesised that the difference between IA- and CA-based risk calculation will be more pronounced when there is higher variability in RQ among substances. The reason is that higher among-substance variability will reduce the IA method’s multiplication component

(Π_1_^n^(1-P)) more than it will reduce the CA method’s addition component (Σ_1_^n^ RQ). Thus, with higher variability, the IA-based mixture risk (1-(Π_1_^n^(1-P)) can be expected to increase more than the CA-based risk (Table 1). This suggests that IA-based mixture risk calculation can become more protective in situations where several substances contribute to mixture risk to various degrees, compared to situations where a few substances dominate the mixture risk. However, such hypotheses remain to be tested for different combinations of substances, groups and risk levels.

### 5.3. Conclusions and outlook

This paper has introduced and explored a novel probabilistic approach to mixture risk characterisation, which combines the traditional concepts of CA (concentration addition) and IA (independent action) in a multi-level Bayesian network model. This BN serves as an emulation model for a high-resolution stochastic simulation model for chemical exposure (here referred to as the ENCORE exposure model), and applies risk quotients (RQ) as a simple metric for single-substance risk. The BN has been applied to a pilot study, representing 15 pesticides of concern in five river basin districts across Belgium. The probabilistic mixture risk characterisations obtained from the three methods showed small, but consistent differences, in accordance with other relevant studies: CA-based risk being slightly higher than IA-based risk, with the combined CA+IA-based risk as a compromise. Furthermore, this study has demonstrated that sensitivity analysis of the BN can be used for efficient and robust ranking of risk-driving substances as well as substance groups; this procedure will account for both signal strength and uncertainties built into the model. In conclusion, our results suggest that this type of multi-level BN for can be a useful tool for assessment and management of mixture risk, notably, for comparing mixture risk across different scenarios, and for prioritisation of the main risk drivers.

Further refinement of this approach should focus on the model resolution, such as the discretisation of continuous variables (concentrations and RQ), and the spatial units for risk assessment. The extensive amount of output that will be made available from the ENCORE exposure model, (simulated concentrations of pesticides, pharmaceuticals and industrial substances for 42 countries) will provide many opportunities for further developments, extensions, applications and evaluations of the probabilistic mixture risk approach presented here.

## List of Supplementary Material

### Background information

File 1: ENCORE_MixRisk_SM_01_Spatial.xlsx. Content: Information on Subcatchments and River Basin Districts.

File 2: ENCORE_MixRisk_SM_02_PEC_values.zip (.cvs, zipped). Content: Simulated concentrations (Predicted Environmental Concentrations).

### Data processing (of PEC)

File 3: ENCORE_MixRisk_SM_03_PEC_figures.zip (.png, zipped). Content: Histograms with fitted statistical distributions, organised in 75 figures (15 substances x 5 RBDs) x 12 plots (months).

File 4: ENCORE_MixRisk_SM_04_PEC_parameters.xlsx. Content: Parameters of fitted statistical mixture distributions for PECs.

### BN model specification

File 5: ENCORE_MixRisk_SM_05_BN_code.txt Content: Complete model code for the BN.

File 6: ENCORE_MixRisk_SM_06_BN_documentation.zip (.html and .png, zipped). Content: System-generated model documentation for selected submodels.

File 7: ENCORE_MixRisk_SM_07_BN_user_interface.pdf. Content: Display of the online user interface to the BN.

File 8: ENCORE_MixRisk_SM_08_BN_discretisation.xlsx. Content: Specification of node states.

File 9: ENCORE_MixRisk_SM_09_BN_expressions.xlsx. Content: Expressions used for generating conditional probability tables.

### BN model output

File 10: ENCORE_MixRisk_SM_10_BN_output_tables.xlsx. Content: BN predictions (posterior probabilities and other derived values) for selected scenarios and nodes.

File 11: ENCORE_MixRisk_SM_11_BN_output_figures.zip. Content: Figures displaying BN predictions: (i) substance-level RQ probability distributions and (ii) mixture-level risk characterisations, for all five RBDs.

File 12: ENCORE_MixRisk_SM_12_BN_sensitivities.xlsx. Content: Tables with sensitivity of all mixture risk nodes (SumSumRQ, AnyRQ, and Any_SumRQ): to all substances (sheet “Substances”) and to all groups (sheet “Groups”).

## Notes

### Competing Interest Statement

The authors have declared no competing interest.

### Funding

- CEFIC-LRI: Project ECO-66

