## Supplemental Files 1-12 for "Chemical Mixtures in River Basins: Combining Additive and Independent Effects with Probabilistic Risk Modelling": ENCORE_MixRisk_SM_07_BN_user_interface.pdf

### A multi-level Bayesian network for predicting probabilistic risk of chemical mixtures

By: Jannicke Moe (Norwegian Institute for Water Research)
WWW: Anders L Madsen, Mark Christiansen (HUGIN EXPERT)

First version: June 2025
Latest update: May 2026

#### Chemical mixtures

**Mixtures** of chemical pollutants in the environment pose threats for aquatic ecosystems globally. There are endless possible combinations of substances in the environment, surpassing any available mixture toxicity data. **Environmental risk assessment (ERA)** for chemical mixture therefore requires pragmatic and robust computational approaches. **Bayesian network (BN)** methodology offers new opportunities for probabilistic risk characterisation for mixtures.

The two main mixture toxicity concepts are **Concentration Addition (CA)** and **Independent Action (IA)**. CA typically predicts slightly higher mixture toxicity than IA, while a combination of CA and IA can give more accurate predictions. Traditionally, the CA method has been more commonly used because of the simpler calculation and the more protective outcome. However, the IA method can more easily be implemented in a probabilistic model.

#### Probabilistic risk calculation

The **ENCORE Mixture BN Pilot** is a **multi-level** probabilistic model developed for **combining the CA and IA** methods, implemented as an OOBN (Object-oriented Bayesian Network). The risk characterisation for a substance is based on a **Risk Quotient (RQ)**, here calculated as the ratio **RQ = PEC/HC5** ( = Predicted Environmental Concentration / Hazardous concentration to 5% of the species)

**Environmental risk** is here defined as the probability (P) of RQ exceeding a given threshold, P(RQ > Threshold_RQ).

A standard threshold for screening-level ERA is Threshold_RQ = 1. Here, the user can select alternative thresholds from 0.001 to 100). For example, Threshold_RQ = 0.1 can represent an early warning situation.

#### How to run the model

The BN can be run by setting evidence for input variables:

- Spatial: Select a value either for River Basin District or for Region
- Temporal: Select a value either for Month or for Season
- RQ threshold: Select a value for each level (RQ, SumRQ, SumSumRQ); can be identical or different

Further details about the approach can be found at the bottom of the page.

#### Inputs:

##### Scenarios

###### Spatial and temporal settings

River Basin District -- select --

Region -- select --

Month -- select --

Season -- select --

###### Exceedance thresholds for Risk Quotients

#### Outputs:

##### Substance level

###### Fungicides

Probability of RQ > Threshold_RQ

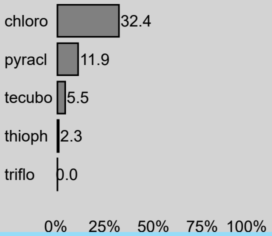


###### Herbicides

#### Outputs:

##### Group level

###### "Concentration Addition" (CA):

Probability of SumRQ > Threshold_SumRQ

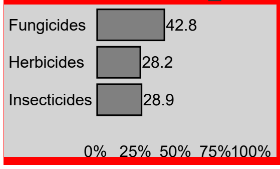


###### "Independent Action" (IA):

#### Outputs:

##### Mixture level

#### "CA":

Probability of SumSumRQ > Threshold_SumSumRQ

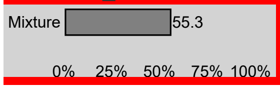


#### "IA":

Probability of one or more RQ > Threshold_RQ

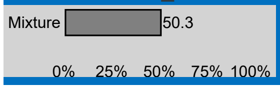

threshold RQ (substance) -- select --
threshold SumRQ (group) -- select --
threshold SumSumRQ -- select --

Probability of
RQ > Threshold_RQ

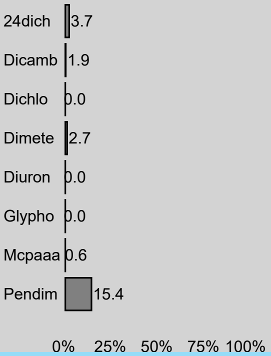


Probability of one or more
RQ > Threshold_RQ

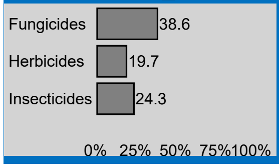


**"CA + IA":**

Probability of one or more
SumRQ > Threshold_SumRQ

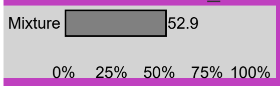


**Insecticides**

Probability of
RQ > Threshold_RQ

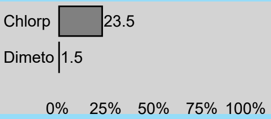


Reset

### Description of pilot study

Spatial extent: Belgium
Spatial resolution: River Basin District (5); Region (2)
Temporal extent: Year 2012
Spatial resolution: Month (12): Season (4)
Chemical substances: 15 pesticides (Chlorothalonil (chloro), Pyraclostrobin (pyracl), Tebuconazole (tebuco), Thiophanate-methyl (thioph), Trifloxystrobin (triflo), 2,4-Dichlorophenoxyacetic acid (24dich), Dicamba (dicamb), Dichlorprop (dichlo), Dimethenamid(-p) (dimete), Diuron (diuron), Glyphosate (glypho), MCPA (mcpaaa), Pendimethalin (pendim), Chlorpyrifos (chlorp), Dimethoate (dimeto))
Chemical groups: 3 groups (Fungicides, Herbicides, Insecticides)

Predicted environmental concentrations are obtained from the ENCORE fate model, a process-based model for simulating chemical concentrations in subcatchments of rivers across Europe.

### Mixture risk calculation with the BN

The BN calculates environmental risk of chemical substances at three levels: substance, substance group, and mixture. The two calculation methods, "Concentration Addition" (CA) and "Independent Action" (IA), are generalisations of the concepts used in ecotoxicology.

1. Substance level: calculation of risk for each substance
   Risk = P(RQ > Threshold_RQ)
2. Group level: calculation of risk for multiple substances within a group
   **CA:**
   SumRQ = $RQ_1 + RQ_2 + ... + RQ_n$
   Risk = P(SumRQ > Threshold_SumRQ)
   **IA:**
   Risk = OR (($RQ_1$ > Threshold_RQ), ($RQ_2$ > Threshold_RQ), ..., ($RQ_n$ > Threshold_RQ))
   = 1 - (1-P($RQ_1$ > Threshold_RQ) × (1-P($RQ_2$ > Threshold_RQ) × ... × (1-P($RQ_n$ > Threshold_RQ))

3. Mixture level: calculation of risk for multiple groups
   **CA:**
   SumSumRQ = $SumRQ_1 + SumRQ_2 + SumRQ_3$
   Risk = P(SumSumRQ > Threshold_SumSumRQ)
   **IA:**
   Risk = OR (($RQ_1$ > Threshold_RQ), ($RQ_2$ > Threshold_RQ), ..., ($RQ_{15}$ > Threshold_RQ))
   **CA + IA (=CA within groups + IA across groups):**
   Risk = OR (($SumRQ_1$ > Threshold_SumRQ), ($SumRQ_2$ > Threshold_SumRQ), ($SumRQ_3$ > Threshold_SumRQ))

**References**

https://www.niva.no/en/projects/encore

Mentzel et al. 2026. ...[in press]

Moe et al. 2026. Chemical Mixtures in European River Basins: A Probabilistic Risk Model for Integrating Additive and Independent Effects. Poster presentation, SETAC Europe 2026.

https://setac.confex.com/setac/europe2026/meetingapp.cgi/Paper/32674

**Acknowledgements**

The project ENCORE (Environmental CO-exposure and Risk Estimation) is sponsored by the European Chemical Industry Council (CEFIC) as part of the Long Range Initiative (LRI) under project ECO-66 *Next generation risk assessment methods supporting the identification of environmental co-exposures of potential concern*.

**Disclaimer**

HUGIN EXPERT A/S takes no responsibility whatsoever for examples and information in examples published on this web site. ALL EXAMPLES ARE FOR DEMONSTRATION PURPOSES ONLY.

---

© HUGIN EXPERT A/S 2025
